# Prior context and individual alpha frequency (IAF) influence predictive processing during language comprehension

**DOI:** 10.1101/2023.05.08.539915

**Authors:** Sophie Jano, Zachariah Cross, Alex Chatburn, Matthias Schlesewsky, Ina Bornkessel-Schlesewsky

**Affiliations:** Cognitive Neuroscience Laboratory, University of South Australia; Feinberg School of Medicine, Northwestern University

## Abstract

The extent to which the brain predicts upcoming information during language processing remains controversial. To shed light on this debate, the present study reanalysed Nieuwland and colleagues’ (2018) replication of DeLong et al. (2005). Participants (*N* = 356) viewed sentences containing articles and nouns of varying predictability, while their electroencephalogram was recorded. We measured event-related potentials (ERPs) preceding the critical words (namely the semantic prediction potential; SPP), in conjunction with post-word N400 patterns and individual neural metrics. ERP activity was compared with two measures of word predictability: cloze probability and lexical surprisal. In contrast to prior literature, SPP amplitudes did not increase as cloze probability increased, suggesting that the component may not reflect prediction during natural language processing. Initial N400 results at the article provided evidence against phonological prediction in language, in line with Nieuwland and colleagues’ findings. Strikingly however, when the surprisal of the prior words in the sentence was included in the analysis, increases in article surprisal were associated with increased N400 amplitudes, consistent with prediction accounts. This relationship between surprisal and N400 amplitude was not observed when the surprisal of the two prior words was low, suggesting that expectation violations at the article may be overlooked under highly predictable conditions. Individual alpha frequency (IAF) also modulated the relationship between article surprisal and the N400, emphasising the importance of individual neural factors for prediction. The present study extends upon existing neurocognitive models of language and prediction more generally, by illuminating the flexible and subject-specific nature of predictive processing.

## Introduction

The question of how the brain converts sensory information into abstract meaning has led to the proposal of prediction as a core mechanism underlying perception (Bar, 2011; Friston, 2005, 2010; Friston & Kiebel, 2009; Press et al., 2020). This is the major assumption of the predictive coding theory, which suggests that prior experiences are used to form generative models (Friston, 2005; Friston & Kiebel, 2009). These models are applied to predict incoming sensory inputs, thus reducing processing effort and facilitating environmental adaptation (Clark, 2013; Friston, 2010; Hohwy, 2013). Although prediction is largely accepted as an explanatory mechanism within numerous cognitive domains (Apps & Tsakiris, 2014; Horga et al., 2014; Koster-Hale & Saxe, 2013), a longstanding debate remains within the cognitive neuroscience of language between prediction-based and integrationist theories of language processing (cf., for example, Ferreira & Chantavarin, 2018). Prediction theories propose that the brain anticipates upcoming linguistic content in a top-down manner, whilst integrationist theories argue that language comprehension proceeds via the bottom-up integration of linguistic units with the prior context (Kuperberg & Jaeger, 2016; Mantegna et al., 2019; Van Petten & Luka, 2012). Although seemingly specific to language, this debate could have important implications for existing understandings of prediction as a ubiquitous cognitive mechanism (Friston, 2010; Kveraga et al., 2007), particularly if linguistic prediction does not occur. As such, clarifying the role of prediction in language processing could provide insight into whether predictive processing is indeed fundamental to general cognition as has been argued (Friston, 2010; Kveraga et al., 2007).

### Prediction in language as revealed by the N400 component

As prediction and integration in language are notoriously difficult to disentangle empirically, the debate surrounding their role in linguistic processing has hinged prominently on an influential study by DeLong et al. (2005). DeLong and colleagues investigated the N400 event-related potential (ERP), which has been linked to prediction errors in language (e.g., Bornkessel-Schlesewsky & Schlesewsky, 2019; Hodapp & Rabovsky, 2021; Rabovsky et al., 2018). By the nature of its design, their experiment sought to exclude an integration-based explanation of purported prediction effects, by capitalising on the English regularity whereby the indefinite articles ‘a’ and ‘an’ precede words beginning with a consonant, and beginning with a vowel, respectively (e.g., “a kite” or “an airplane”). DeLong et al. (2005) observed larger N400 amplitudes time-locked to the article preceding unpredictable (versus predictable) nouns (e.g., “The day was breezy, so the boy went outside to fly *an* …”; the article “an” rules out the predictable continuation “kite”). As the indefinite articles ‘a/an’ have the same meaning, this observation is difficult to reconcile with integrationist views of the N400, which postulate that N400 differences reflect the ease with which a word’s meaning can be integrated with its preceding context (Van Berkum et al., 1999; Brown & Hagoort, 1993; Chwilla & Kolk, 2000). Moreover, the N400 effect was graded in accordance with how predictable the article word was, suggesting that language comprehension involves the prediction and pre-activation of words down to the highly detailed level of speech sounds. This assumption is highly compatible with predictive coding approaches, where predictions are proposed to activate a representation of the anticipated stimulus at multiple levels of the cortical hierarchy (Rao & Ballard, 1999; Rauss & Born, 2017). Therefore, whilst there are various definitions for prediction (for discussion see Kuperberg & Jaeger, 2016), such N400-article analyses have the potential to provide evidence for the prediction of form-based linguistic attributes.

The investigation by DeLong and colleagues (2005) suggests that prediction serves as a likely mechanism for linguistic processing. This is supported by research which has observed similar ERP effects time-locked to adjectives (Van Berkum et al., 2005) and gender-marked determiners (Wicha et al., 2004) preceding a noun, as well as prior to sign onset in sign language (Hosemann et al., 2013; Krebs et al., 2019). However, other studies have observed differential patterns (for discussion see Ito et al., 2017; Schoknecht et al., 2022), including a large-scale direct replication study by Nieuwland et al. (2018), who analysed data from 356 participants using the same stimuli as in DeLong et al. (2005). In contrast to DeLong and colleagues, Nieuwland and colleagues did not observe a more pronounced N400 following unpredictable articles. Accordingly, they suggested that, although the brain may predict upcoming contextual or semantic information, this may not extend to form-based linguistic characteristics such as speech sounds (Nieuwland et al., 2018). Therefore, the extent of prediction in language is currently unclear and may be subject to significant variability. Consequently, the mixed nature of the literature renders it important to examine other sources of variability that may influence predictive processing.

### The influence of individual neural factors on prediction

An investigation of variability at the individual level may provide insight into linguistic prediction, with such variability capturing effects over and above group-level contrasts in previous studies (e.g., Roehm et al., 2013). Although individual neural differences were not considered by DeLong et al. (2005), nor by Nieuwland et al. (2018), such activity could contribute to understandings of prediction in language, particularly if prediction tendencies differ between individuals. A potential neurobiological marker that may relate to individual differences in prediction is the individual alpha frequency (IAF), the individual peak frequency within the alpha rhythm (Corcoran et al., 2018; Grandy et al., 2013a; Klimesch, 1999). The link between IAF and prediction primarily stems from IAF’s role as a trait-like marker of information processing, where those with a high IAF were shown to have faster reaction times (Klimesch et al., 1996; Surwillo, 1961, 1963) and improved discrimination of rapidly presented stimuli (Samaha & Postle, 2015). Enhanced information processing in high versus low IAF individuals may therefore translate to a stronger reliance on bottom-up perceptual information as opposed to a top-down model, resulting in more frequent model updating (Kurthen et al., 2020; however, see Bornkessel-Schlesewsky et al., 2022 for a stronger tendency to adjust predictions in low IAF individuals). As such, IAF may influence predictive processing and the extent to which sensory information is used to adjust predictions.

Despite periodic oscillatory components such as IAF traditionally being linked to cognitive processing, the importance of considering the aperiodic noise-like activity (particularly the 1/f slope of the distribution) that underlies the EEG spectra has been recently emphasised in the literature (Cross et al., 2022; Demuru & Fraschini, 2020; Dziego et al., 2023; Immink et al., 2021; Merkin et al., 2023; Ouyang et al., 2020; Voytek et al., 2015; Voytek & Knight, 2015). Although prior literature has ignored or dismissed this activity as noise (Donoghue et al., 2020), the 1/f component has been shown to predict information processing speeds (Ouyang et al., 2020), and to mediate the relationship between age and cognitive functioning (Pathania et al., 2022; Voytek et al., 2015), suggesting that it may be relevant for cognition. Regarding prediction in language, steeper 1/f slopes have been related to prediction error processing (Cross et al., 2022), stronger N400 effects (Dave et al., 2018), and to a greater tendency to update a predictive model (Bornkessel-Schlesewsky et al., 2022). Therefore, by analysing the role of the 1/f slope in the Nieuwland et al. (2018) dataset, it may be possible to capture the individual variability relating to prediction, contributing insight to the extent of prediction in language and the role of inter-subject variability in predictive processing.

### Pre-stimulus event-related potentials (ERPs) may provide insight into linguistic prediction

Whilst an analysis of individual factors may shed light on prediction, pre-stimulus ERP activity may also probe predictive processing more directly than post-word N400 patterns. Pre-stimulus ERPs have been proposed to reflect the transient, time-locked activity associated with anticipation (Grisoni et al., 2016, 2017, 2019, 2021). Such components are generally negative-going deflections with frontal topographies and larger amplitudes preceding predictable information (Grisoni et al., 2021; León-Cabrera et al., 2019). Of note is the semantic prediction potential (SPP), which exhibits greater amplitudes in the interval preceding predictable versus unpredictable critical words, whilst demonstrating topographical variance in accordance with the meaning of each word (Grisoni et al., 2017, 2021). This provides evidence not only for prediction but for the pre-activation of the semantic characteristics of a word before it arises (Grisoni et al., 2017, 2021). Such correlates also strongly resemble the stimulus preceding negativity (SPN) ERP (León-Cabrera et al., 2019), a component associated with the subjective appraisal of prediction accuracy (Ono et al., 2021), and with feedback anticipation during reinforcement learning (Brunia et al., 2011; Morís et al., 2013). As such, whilst the absence of an N400 effect at the article (as observed in Nieuwland and colleagues’ original analysis) was considered as evidence against the prediction of detailed linguistic features, a relationship between SPP amplitude and word predictability in the pre-stimulus window may provide alternative support for prediction in language, at least at the semantic level. Further, the SPP has been investigated in contexts alongside the N400 component, with high (versus low) constraint sentences eliciting larger pre-word SPP amplitudes and smaller N400 amplitudes (Grisoni et al., 2021). This inverse relationship between the N400 and the SPP may be due to the predictable nature of the words in the high-constraint sentences, which likely facilitates strong predictions (as evidenced by larger SPPs) whilst eliciting fewer prediction violations (potentially reflected in decreased N400 amplitudes; Grisoni et al., 2021). Consequently, this relationship between the SPP and the N400, and the correlations between the SPP and word predictability, highlight the potential utility of the component for understandings of prediction in language.

### The present study

The current study aimed to investigate the extent of prediction in language and the role of individual neural differences in predictive processing. To do so, we analysed the event-related activity preceding the critical words in the Nieuwland et al. (2018) data (i.e., the SPP), in conjunction with the 1/f aperiodic slope and IAF. Such an analysis has the potential to shed light on the discrepancies in findings between Delong et al. (2005) and Nieuwland et al. (2018), while concomitantly deepening existing understandings of prediction as a general cognitive mechanism (Friston, 2010). It was hypothesised that increases in article and noun predictability would be associated with greater pre-stimulus SPP amplitudes, and that this would be modulated by IAF and the 1/f slope. Additionally, although not the original focus of the study, we report analyses of the N400 component to maintain consistency with Nieuwland and colleagues’ investigation. After initial plots revealed differences in ERP activity at earlier stages of the sentence, we also conducted exploratory analyses including prior word surprisal, shedding light on the role of prior context in prediction.

## Methods

### Original study

The present study involved a reanalysis of Nieuwland et al. (2018), which included 356 participants (females = 222, mean age = 19.8 years) who were all right-handed native English speakers. Participants were recruited across nine different laboratories and the sample size was chosen to ensure that each laboratory’s sample equated approximately to that of DeLong et al. (2005), who included 32 participants in total (Nieuwland et al., 2018). Participants provided informed consent for the use of their data, and ethical approval for each laboratory was sought in accordance with the requirements of the specific institution (Nieuwland et al., 2018). Cloze probability (the probability that a word will be chosen as the likely continuation based on the prior sentence context; Taylor, 1953) was used as an index of stimulus predictability. The materials consisted of 80 sentences with two cloze probability continuations: a more-expected article/noun combination (e.g., “…the boy went outside to fly *a kite*”) and a less-expected article/noun combination (e.g., “…the boy went outside to fly *an airplane*”). Words within each sentence were presented for 200ms sequentially on a computer screen and separated by a 300ms interstimulus interval during which participants viewed a blank screen. Following 25% of sentences, participants answered a yes/no comprehension question. EEG was recorded throughout the study to enable an analysis of the event-related activity in response to articles and nouns. For a detailed description of the study protocol and materials, the reader is directed to Nieuwland et al. (2018).

### Participants

Data for 356 participants were originally obtained from the Open Science Framework (OSF) repository (see https://osf.io/eyzaq/). Five participants were removed prior to the BIDS conversion (see *Data processing* section) due to incomplete files, answering questions randomly or due to being a non-native English speaker. As such, the BIDS conversion was completed for 351 participants. Subsequent pre-processing and epoching led to the removal of 18 more participants (ten due to trigger issues that could not be resolved, and eight participants due to excessive signal noise). As such, data for 333 participants were epoched, with four participants being excluded due to a lack of trials.

Seventeen participants were also identified as having an insufficient number of epochs for the calculation of the aperiodic (1/f) slope and IAF (less than 16 trials). As all analyses included individual factors, these participants were excluded from the final sample, taking the final sample size to 312 (females = 194, mean age = 22.4 years). See *Figure 1* for a schematic of participants at each stage of the data analysis pipeline.

**Figure 1.**
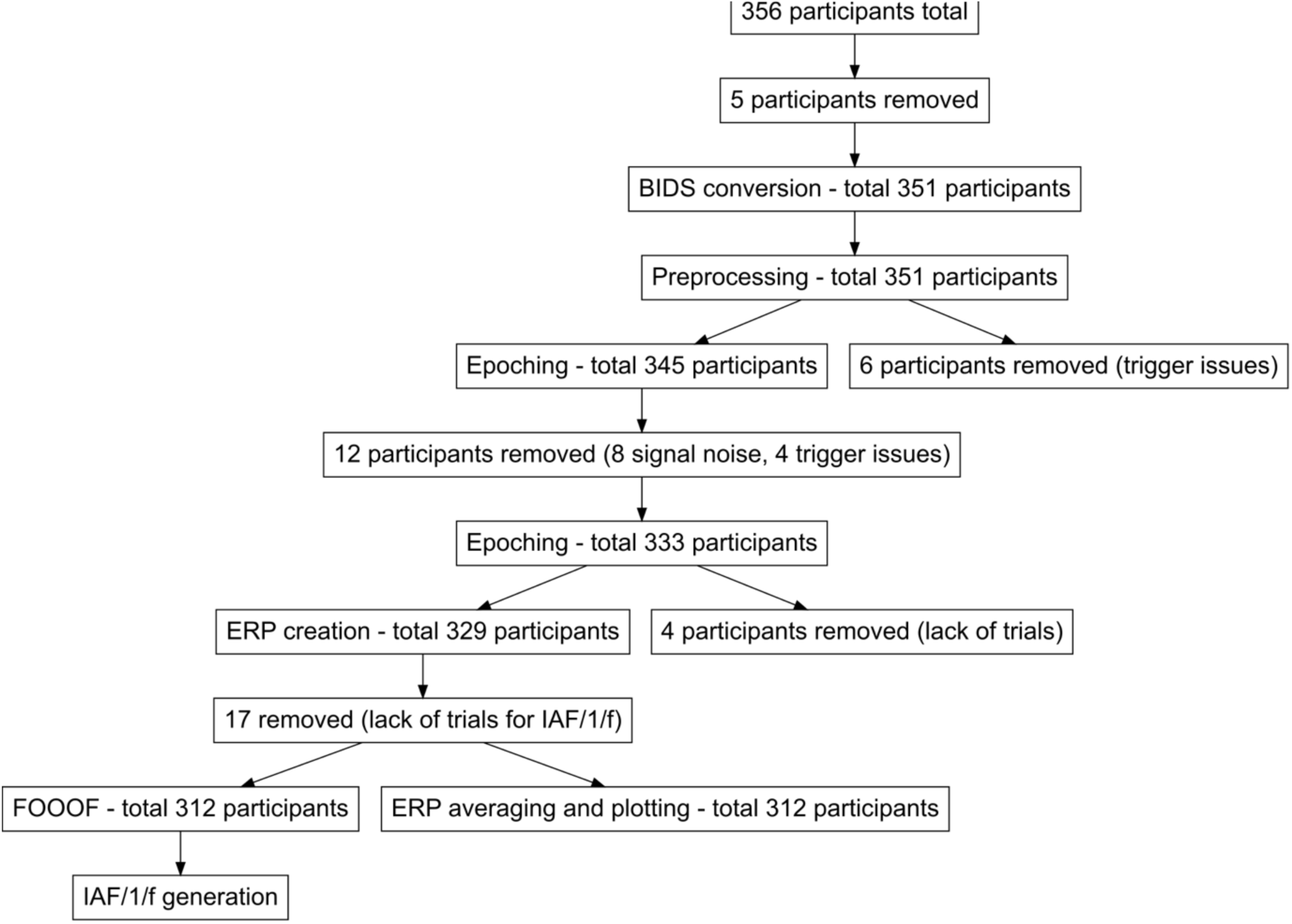
Bayesian sensitivity analysis of the effect of article cloze probability on SPP amplitude preceding the articles. Bayes factors (log transformed) are presented on the Y axis, which are plotted according to each prior standard deviation for the predictor (β; cloze probability).

### Data Processing

Data were first converted to the Brain Imaging Data Structure (BIDS) using the MNE-BIDS package (Appelhoff et al., 2019) in *MNE-python* version 3.8.8 (Gramfort et al., 2013), to account for file structure variations as a result of the differing EEG systems used across the laboratories. As files with the .cnt extension were unsupported by the BIDS package in python (*MNE-BIDS 0.10 Documentation*, 2022), such files were converted to the Brainvision Core Data format (Brain Products Gmbh) prior to the BIDS conversion using MATLAB (MathWorks, Natick, Massachusetts, USA). As most raw files encompassed both the experimental session and the control session, each file was then cropped to exclude the control session, as this was not of interest to the present analysis.

Data were then re-sampled to either 250Hz or 256Hz, depending on the original sampling rate, which ranged from 250Hz to 2048Hz between laboratories. Resampling occurred at an equal interval to avoid data loss when epoching (Gramfort et al., 2013), hence the two sampling frequencies. Therefore, the final sample consisted of 184 participants with data sampled at 250Hz, and 145 participants with sampling rates of 256Hz. This is not expected to have affected the single-trial ERP analyses, as samples were averaged across time intervals of interest and slight discrepancies in the number of samples per window should not introduce significant variability. Following resampling, the EEG data were re-referenced to linked mastoids, while EOGs were set as Fp1 and Fp2 across laboratories. For laboratories that used the alphabetized Biosemi system (BioSemi, 2019), EOG channels were those closest to Fp1 and Fp2 (i.e., C29 and C16, respectively). The data were filtered using a bandpass filter that ranged from 0.1 to 30Hz.

To conduct an independent component analysis (ICA), temporary epochs around events of interest were first generated. Rejection thresholds were then computed from the epochs using the *Autoreject* package in MNE-python (Jas et al, 2016), to be included in the ICA for the detection and removal of ocular artefacts. This procedure is argued to lead to more precision in the ICA, as thresholds are tailored specifically depending on the activity in segments of interest (Jas et al., 2016). The ICA was then fit to a 1Hz high pass filtered copy of the temporary epochs, to minimise the influence of noise from the inter-trial intervals. Subsequently, the ICA was applied to the raw data (See *Table 1* for average number of trials per condition). Following this, channels marked as bad were interpolated using spherical splines and the raw data were epoched around events of interest into segments ranging from -800ms to 1000ms (1.8 seconds total). The *Autoreject* function (Jas et al, 2016) was then applied to repair or remove bad epochs (average percentage of epochs rejected across the final 312 participants = 8.96%).

**Table 1.** Demonstrating the mean number of trials per condition that were entered into the ICA, with standard deviations (SD) in brackets.

| Expected articles | Unexpected articles | Expected nouns | Unexpected nouns | Article-1* |
| --- | --- | --- | --- | --- |
| 39.97 (0.29) | 39.98 (0.20) | 39.97 (0.30) | 38.96 (1.01) | 79.98 (0.15) |

*\*The article-1 window (-300 to 0ms preceding the word before the article) was used as a baseline period.*

### ERP analysis

Event-related activity was extracted within the SPP and N400 time windows on a trial-by-trial basis. The SPP window ranged from -300 to 0ms prior to the article and noun words, to encompass the length of the interstimulus intervals. The N400 time window ranged from 200ms to 500ms post-stimulus onset, in line with Nieuwland et al. (2018). To control for differences in neural activity unrelated to the experimental manipulation, epochs were also generated around the word preceding the article. The inter-stimulus interval (−300 to 0ms) preceding the word before the article (the article-1 word) was chosen as a baseline period due to its temporal proximity to the trials of interest, and because the SPP activity preceding the articles and nouns could not be used since it was the focus of the present investigation. This pre-article-1 word baseline window was not used for traditional baseline correction, but rather the mean amplitude across this window was entered as a covariate into the statistical models of SPP effects involving both the article and the noun that followed (in line with Alday, 2019). For N400 analyses at the article, activity in the pre-article window itself (−300 to 0ms preceding the article) was used as the “baseline” period, whilst for noun N400 analyses, activity in the pre-noun window (−300 to 0ms preceding the noun) was used. This activity was similarly included as a covariate for statistical modelling, and according to Alday (2019), does not need to be further interpreted once controlled for (i.e., it serves as a covariate of no interest).

To generate ERPs, *MNE-Python* (Gramfort et al., 2013) was used to calculate the mean ERP amplitude across time windows of interest. Due to differing EEG systems, one laboratory lacked channels that directly corresponded to a common 10-20 channel shared by the other laboratories. As such, a region-based approach was taken whereby SPP event-related activity was analysed over frontal regions, in line with previous research (León-Cabrera et al., 2017, 2019; Morís et al., 2013). For N400 analyses, we included the channels that were used in the original study (Cz, C3, C4, Pz, P3, and P4). However, the laboratory that used the alphabetized system did not have channels that directly corresponded to P3 and P4, and thus the closest channels were used (A7 and B4, respectively). ERP grand average plots were created using *ggplot2* in *R version 4.1.1* (Wickham, 2016). Due to the differing sampling intervals, the timing of the samples for four laboratories was adjusted by a maximum of four milliseconds to match the timing of the other laboratories. This was done for visualisation purposes only. Additionally, for ease of visualisation, channel-specific ERP plots include data from the alphabetized Biosemi channel that was closest to the shared 10-20 channel nomenclature.

### Individual alpha frequency and aperiodic 1/f slope estimation

Individual alpha frequency (IAF) and the aperiodic 1/f slope were calculated at the individual level using the *Fitting Oscillations & one-over f (FOOOF)* package (version 1.0.0) in *MNE-python* (Gramfort et al., 2013). Epochs were first generated around the between sentence intervals as during this time, participants should not have been actively engaged in the task (in exception is the 25% of trials that included a comprehension question, which were removed), and thus task-related effects were likely reduced. Epochs ranged from 200ms after the final word (when the presentation of the final word had ended), to 2.2 seconds, to ensure a sufficient number of samples to obtain reliable estimates. As the between-sentence intervals were self-paced, trials in which the intervals were shorter than 2.2 seconds were removed from the analysis. Similarly to the treatment of task-related epochs, the *Autoreject* function (Jas et al., 2017) was then applied to the between-sentence epochs to repair or remove bad trials.

Following epoching, the *FOOOF* algorithm was fit to PSD estimates derived from each channel of interest, per epoch (Donoghue et al., 2020). Channels Oz, O1, and O2 were selected, as alpha activity is prominent over occipital regions (Grandy et al., 2013b), and as prior research suggests that the relationship between 1/f and cognitive processing may be similar across the scalp (Ouyang et al., 2020). For the laboratory with the alphabetized system, the channels closest to Oz, O1 and O2 were used (A23, A15 and A28, respectively). Parameterization of the power spectra occurred across a frequency range of 1-30Hz. Settings for the *FOOOF* algorithm included a peak width limit of 2 to 8, a maximum peak number of four, the minimum peak height as 0.0, a peak threshold of 2.0, and a fixed aperiodic mode. To obtain individual metrics, activity across the three channels was averaged across all epochs, resulting in single-subject slope and offset values. IAF estimates were calculated by selecting peaks within the 7-13Hz range and averaging over the activity across each channel and epoch to obtain a single IAF value per participant.

### Statistical analysis

Statistical model structures for the primary analyses are presented in *Table 3.* To examine effects of interest whilst accounting for random effects, linear mixed-effects models (LMMs) were computed using the *Lme4* package in *R version 4.2.2* (Bates et al., 2015). P-values were calculated using the *summary* function from the *lmerTest* package, which is based on Satterthwaite’s degrees of freedom (Kuznetsova et al., 2017). The alpha level was set at 0.05 and all predictor variables were scaled and centred. Note that for the main analyses, IAF and the 1/f slope were included in the same models, specifying different interaction terms, as the relationship between the two factors was not a primary interest of the study. The model fits of these combined models also were either better or did not significantly differ from separate models for IAF and 1/f, as measured by the Akaike criterion (See *Appendix 1 – Table 1* for Akaike criterion values).

Primary and exploratory surprisal models (described in the *Exploratory surprisal analyses* section) initially failed to converge with complex random effects structures (i.e., random intercepts of item, subject and channel, and by-item and by-subject random slopes). To simplify the model structures, ERP activity was averaged over channels of interest, allowing for the removal of random intercepts of channel. However, the models either failed to converge, or exhibited singular fits, when random slopes according to all predictors were included. To avoid overfitting, by-subject and by-item random slopes according to the primary predictor of interest (critical word cloze or critical word surprisal) were instead included (for a discussion of model overspecification resulting from maximal LMM structures, see Bates et al., 2018). However, one model (model 1) failed to converge with random slopes, which were omitted. To determine whether the inclusion of random slopes improved goodness of fit, models that converged with random slopes were compared with and without the random slopes using Likelihood ratio tests (LRTs). The LRTs revealed that for all models (excluding Model 3), the inclusion of random slopes significantly improved the model fit (See *Appendix 1 – Table 2* for LRT results). Model 3 also initially only exhibited a non-singular fit with by-item, not by-subject random slopes. However, because the by-item random slopes did not improve the model fit, they were removed.

When investigating the effect of word predictability (operationalized as cloze probability) on the SPP time-locked to articles (Model 1), cloze probability was contrasted between-sentences, as opposed to between expected and unexpected sentence conditions. This is because, prior to the article, the sentence contexts were identical. The likely continuation (the expected article cloze value for that sentence) was taken as the cloze metric for both versions of the sentence (See *Table 2* for an example). In contrast, when measuring SPP effects time-locked to the nouns, the original noun cloze values for both expected and unexpected continuations were applied because the sentences arguably differed once participants had been exposed to the article variation.

**Table 2.** Example sentence demonstrating the cloze probability values (in brackets) for both the unexpected and expected continuations. *Note that prior to the article, the sentence versions are identical, thus justifying the application of the expected article cloze value (82) to all versions of the sentence when analysing pre-stimulus effects time-locked to articles*.

| Sentence beginning | Expected article | Unexpected article | Expected noun | Unexpected noun |
| --- | --- | --- | --- | --- |
| The old wives' tale says that if you want to keep the doctor away then you should eat | an (82) | a (0) | apple (97) | carrot (0) |

**Table 3.** Linear mixed-effects models for primary analyses with outcome variables, covariates, fixed and random effects. Note that IAF and 1/f were included as separate interaction terms, such that in model 1 for example, the interaction of cloze*IAF was examined independently from the interaction of cloze*1/f slope.

| **Model** | **Outcome** | **Covariates** | **Fixed effects** | **Separate interaction terms** | **Random intercepts** | **Random slopes** |
| --- | --- | --- | --- | --- | --- | --- |
| 1 | **Pre-article SPP**<br>amplitude<br>(-300 to 0ms) | Article-1 word ERP<br>amplitude<br>(-300 to 0ms) | Between-sentence<br>article cloze<br>probability | 1/f slope<br>IAF | + Subject<br>+ item |  |
| 2 | **Pre-noun SPP**<br>amplitude<br>(-300 to 0ms) | Article-1 word ERP<br>amplitude<br>(-300 to 0ms) | Noun cloze<br>probability | 1/f slope<br>IAF | + Subject<br>+ item | By-subject<br>and by-item<br>according to<br>cloze |
| 3 | **Post-article N400**<br>amplitude<br>(200 to 500ms) | Pre-article ERP<br>amplitude<br>(-300 to 0ms) | Article cloze<br>probability | 1/f slope<br>IAF | + Subject<br>+ item |  |
| 4 | **Post-noun N400**<br>amplitude<br>(200 to 500ms) | Pre-noun ERP<br>amplitude<br>(-300 to 0ms) | Noun cloze<br>probability | 1/f slope<br>IAF | + Subject<br>+ item | By-subject<br>and by-item<br>according to<br>cloze |

The model effects were plotted using the *ggplot2* package (Wickham, 2016), with 83% confidence intervals to enable visual identification of statistical effects. For plots in which variables are presented dichotomously (e.g., high/low), this distinction was categorised according to the first and third quartiles.

### Exploratory surprisal analyses

Early ERP grand average plots revealed event-related differences according to between-sentence cloze probability at earlier stages of the sentence (See *Figure 2*). This sparked interest regarding the accumulation of predictions over time, and of how the predictability of the prior words in the sentence may influence later predictions. Additional exploratory LMMs were thus included to analyse whether the surprisal of the preceding words interacted with the surprisal of the articles to influence ERP amplitudes time-locked to the articles. The raw surprisal values of the two preceding words (article-2 and article-1) and of the critical articles were included in the statistical models. To maintain consistency with our cloze models, LMMs were also computed investigating the effect of article/noun surprisal alone on ERP amplitude time-locked to the articles and nouns, respectively (See *Table 4* for exploratory model structures). IAF and the 1/f slope were included in all exploratory statistical models, and random effects were structured similarly to the original models. Note that although we ran SPP models with surprisal, the results either did not differ to those of cloze or did not reveal interactions with the surprisal of the critical word. Therefore, for simplicity’s sake, we do not report the results of these models or specify their structures in *Table 4*, although the model outputs are presented in *Appendix 3*. Additionally, because an N400 effect at the noun could not directly provide evidence for phonological prediction in the same manner as effects at the article (where a relationship between N400 amplitude and word predictability may index prediction for the upcoming noun), we did not deem it necessary to examine prior word surprisal in this context. Rather, analyses at the noun were primarily included as a sanity check and for completeness. Therefore, to reduce complexity, we do not report N400 effects at the noun with prior word surprisal. Finally, Pearson correlations were calculated to examine the relationship between cloze probability and surprisal for the two respective word types.

**Figure 2.**
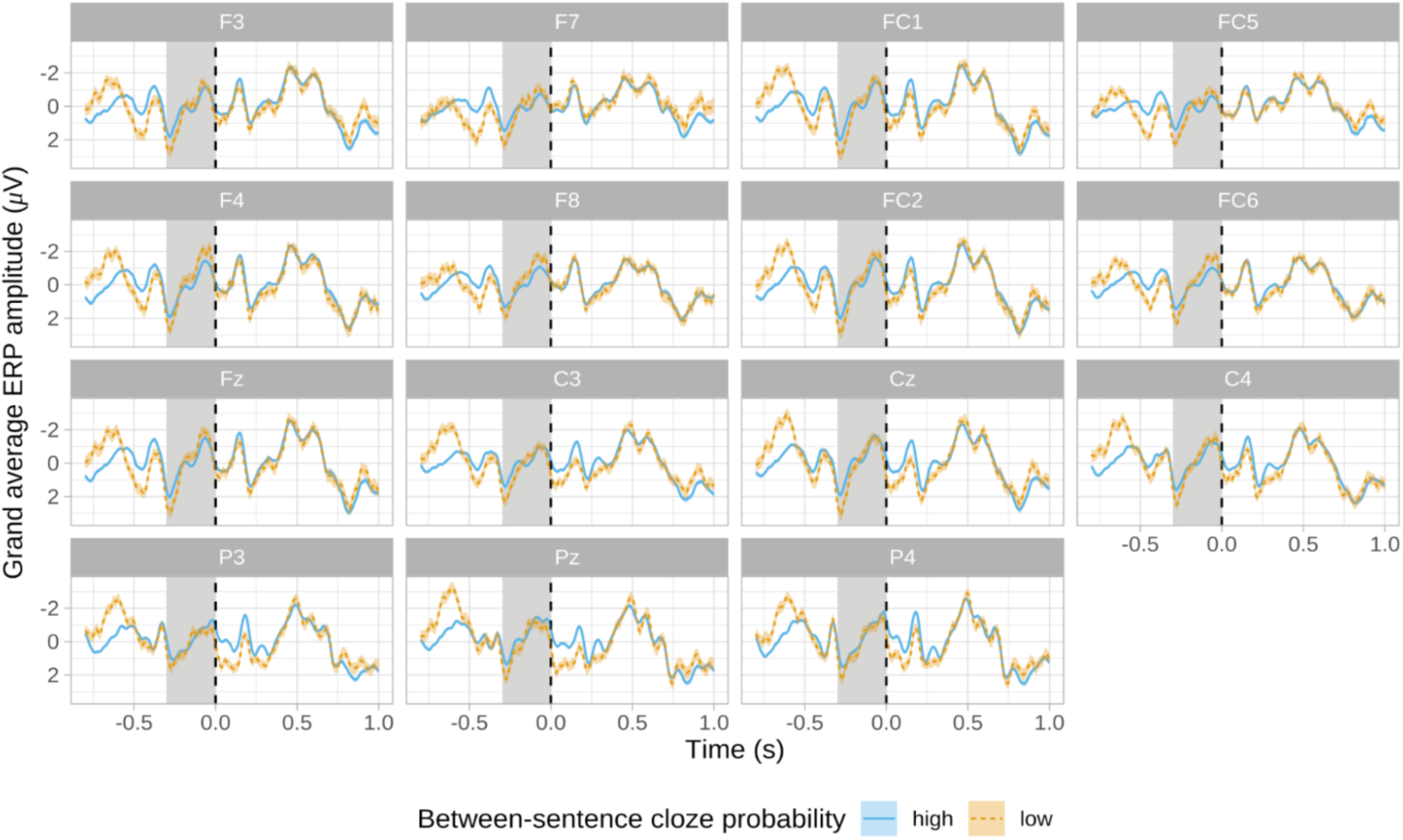
Bayesian sensitivity analysis of the effect of noun cloze probability on SPP amplitude preceding the nouns. Bayes factors (log transformed) are presented on the Y axis, which are plotted according to each prior standard deviation for the predictor (β; cloze probability).

**Table 4.** Exploratory linear mixed-effects model structures with outcome variables, covariates, fixed and random effects.

| **Model** | **Outcome** | **Covariates** | **Fixed effects** | **Separate interaction terms** | **Random intercepts** | **Random slopes** |
| --- | --- | --- | --- | --- | --- | --- |
| 5 | **Post-article**<br>N400<br>amplitude<br>(200 to<br>500ms) | Pre-article<br>ERP<br>amplitude | Article<br>surprisal | IAF<br>1/f | + Subject<br>+ item | By-subject<br>and by-item<br>according to<br>article<br>surprisal |
| 6a | **Post-article**<br>N400<br>amplitude<br>(200 to<br>500ms) | Pre-article<br>ERP<br>amplitude | Article<br>surprisal<br>Article-1<br>surprisal<br>Article-2<br>surprisal<br>1/f |  | + Subject<br>+ item | By-subject<br>and by-item<br>according to<br>article<br>surprisal |
| 6b | **Post-article**<br>N400<br>amplitude<br>(200 to<br>500ms) | Pre-article<br>ERP<br>amplitude | Article<br>surprisal<br>Article-1<br>surprisal<br>Article-2<br>surprisal<br>IAF |  | + Subject<br>+ item | By-subject<br>and by-item<br>according to<br>article<br>surprisal |
| 7 | **Post-noun**<br>N400<br>amplitude<br>(200 to<br>500ms) | Pre-noun<br>ERP<br>amplitude | Noun<br>surprisal | IAF<br>1/f | + Subject<br>+ item | By-subject<br>and by-item<br>according to<br>noun<br>surprisal |

Lexical surprisal was calculated using an open artificial intelligence model known as Generative Pre-Trained Transformer-2 (GPT-2; Radford et al., 2019) using the *Transformers* package in Python (Wolf et al., 2020). Surprisal in a linguistic context is calculated based on the probability of a given word in a linguistic corpus or database, whereby a higher surprisal value corresponds to lower predictability of a word within, for example, a sentence (Frank et al., 2013). GPT-2 is trained on a large corpus of English texts to traditionally predict upcoming words in a sentence, hence its efficacy in calculating surprisal (Radford et al., 2019). This measure was specifically used as the original experiment lacked cloze probability values of the prior words (Nieuwland et al., 2018).

## Results

Statistical model outputs are presented in *Appendix 2*.

### SPP effects

SPP ERP grand average plots time-locked to articles are shown in *Figure 2,* and ERP effects for nouns are shown in *Figure 3.* It was hypothesized that SPP amplitude directly preceding the articles would become larger (i.e., more negative) as article predictability increased, and that this relationship would be influenced by IAF and the 1/f slope (tested via Model 1). The main effect of between-sentence article cloze probability on pre-article SPP amplitude was nonsignificant (*p* = 0.620, β = 0.06, *SE* = 0.13, 95% CI = -0.18 – 0.31), similarly to the interaction of between-sentence article cloze probability and IAF on pre-article SPP amplitude (*p* = 0.748, β = -0.03, *SE* = 0.08, 95% CI = -0.19 – 0.13). The effect of between-sentence article cloze and 1/f slope on SPP amplitude preceding the articles was also nonsignificant (*p* = 0.720, β = -0.03, *SE* = 0.08, 95% CI = -0.19 – 0.13). Therefore, there were no detectable effects of article cloze probability and individual factors on SPP activity before the articles.

**Figure 3.**
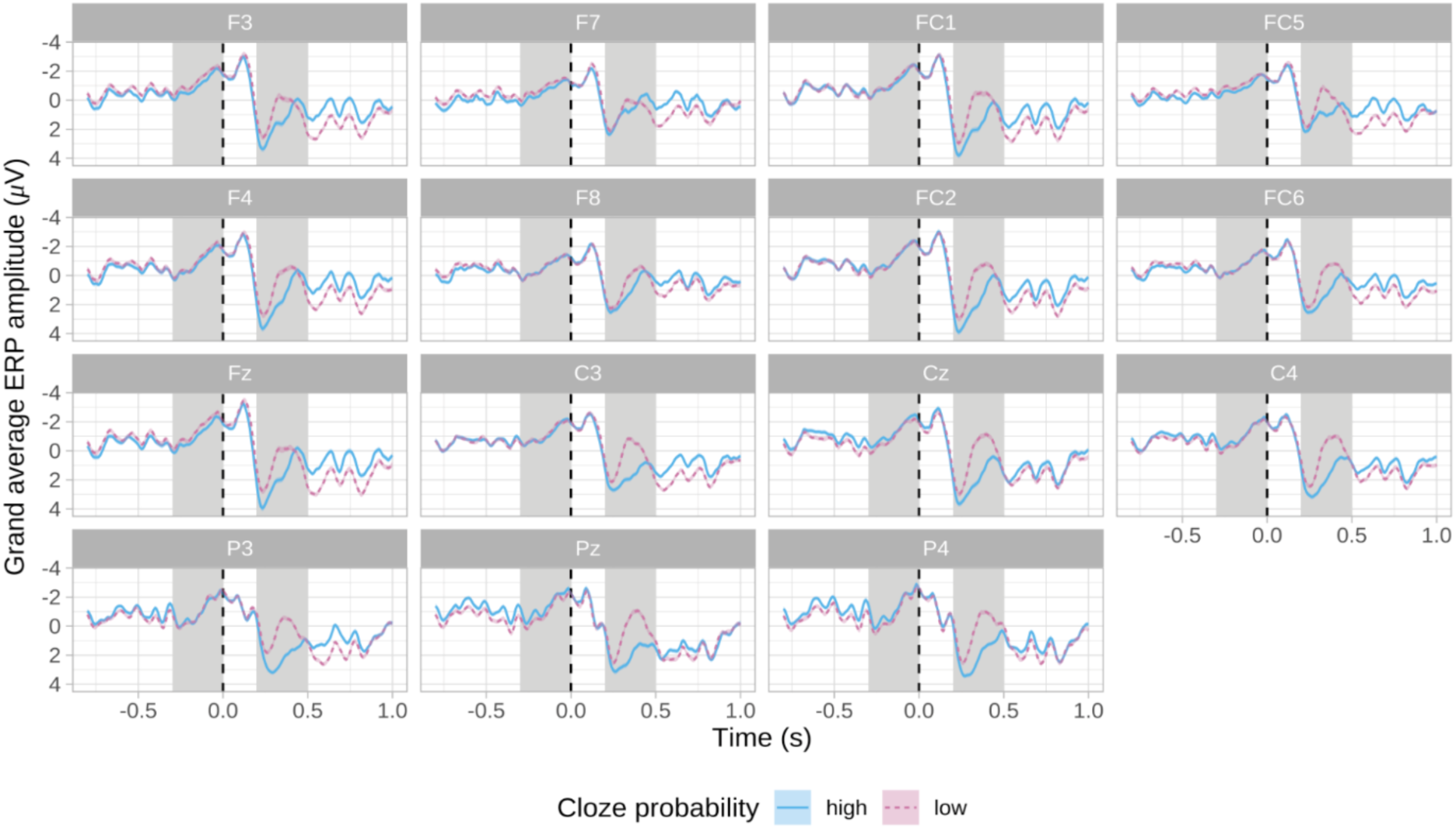
Grand average ERP plots for noun words over channels of interest according to noun cloze probability (*n* = 312). Blue lines represent high noun cloze probability (>50) whilst magenta dashed lines reflect low noun cloze probability (<50). Blue and magenta shading reflects the standard errors for the respective conditions. The SPP window and the N400 window are denoted via grey shading, and the black dashed line depicts the onset of the noun word. Note that the plot depicts non-baseline corrected ERP activity.

Model 2 addressed the hypothesis that increases in noun cloze probability would lead to greater (i.e., more negative) SPP amplitudes directly preceding the nouns, and that this would be modulated by IAF or the aperiodic 1/f slope. The model demonstrated a significant main effect of noun cloze probability on pre-noun SPP amplitude (*p* = 0.038, β = 0.11, *SE* = 0.05, 95% CI = 0.01 – 0.21), such that increases in noun cloze probability were associated with decreased (i.e., less negative) SPP amplitudes preceding the nouns (see *Figure 4*). The interaction between noun cloze and IAF on pre-noun SPP amplitude was non-significant (*p* = 0.220, β = - 0.05, *SE* = 0.04, 95% CI = -0.12 – 0.03), similarly to the effect of noun cloze and 1/f slope (*p* = 0.919, β = -0.00, *SE* = 0.04, 95% CI = -0.08 – 0.07). As such, we did not detect a modulatory effect of individual factors on the relationship between cloze and the SPP.

**Figure 4.**
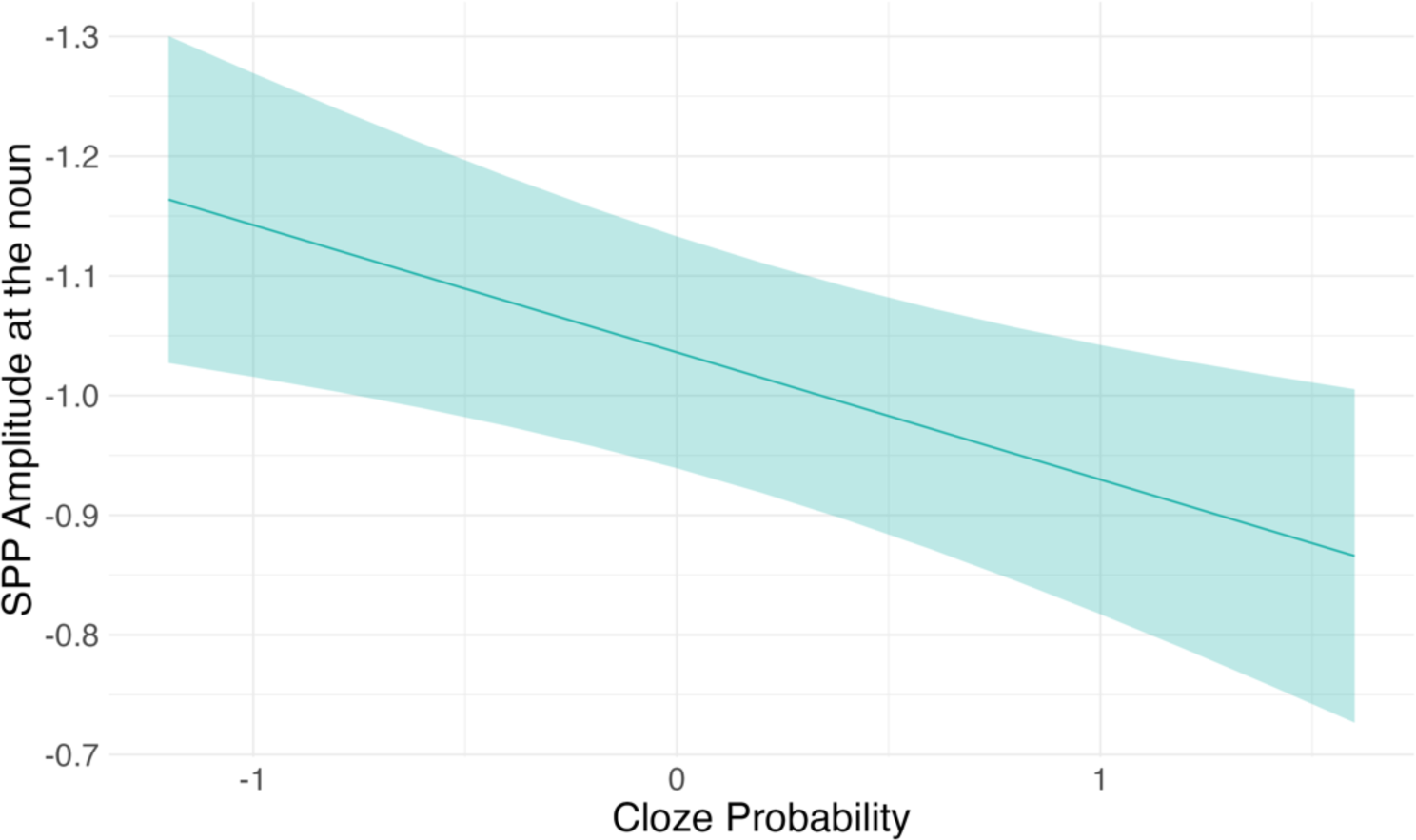
Linear mixed model results depicting the effect of noun cloze probability (scaled and centered) on SPP amplitude time-locked to the nouns (*p* = 0.038, β = 0.11, *SE* = 0.05, 95% CI = 0.01 – 0.21). Lines represent model fit, with shading depicting the 83% confidence intervals.

To further quantify the effects of cloze probability on SPP amplitude, we ran exploratory Bayesian sensitivity analyses, as recommended by an anonymous reviewer (described in *Appendix 4*). Broadly, the results provided additional evidence against an effect of cloze probability on SPP amplitude.

### N400 ERP effects

N400 grand average ERPs time-locked to articles are presented in *Figure 5.* To examine whether individual neural factors influence the relationship between article cloze probability and N400 amplitude directly following the article, Model 3 was fitted. The model revealed a significant effect of article cloze probability on N400 amplitude time-locked to the article (*p* <.001, β = -0.14, *SE =* 0.03, 95% CI = -0.20 – -0.07), such that as article cloze increased, post-article N400 amplitudes increased (i.e., became more negative). The interaction between article cloze and IAF was nonsignificant (*p* = 0.937, β = 0.00, *SE =* 0.03, 95% CI = -0.06– 0.07), as was the interaction between article cloze and 1/f slope (*p* = 0.512, β = -0.02, *SE =* 0.03, 95% CI = -0.09 – 0.04).

**Figure 5.**
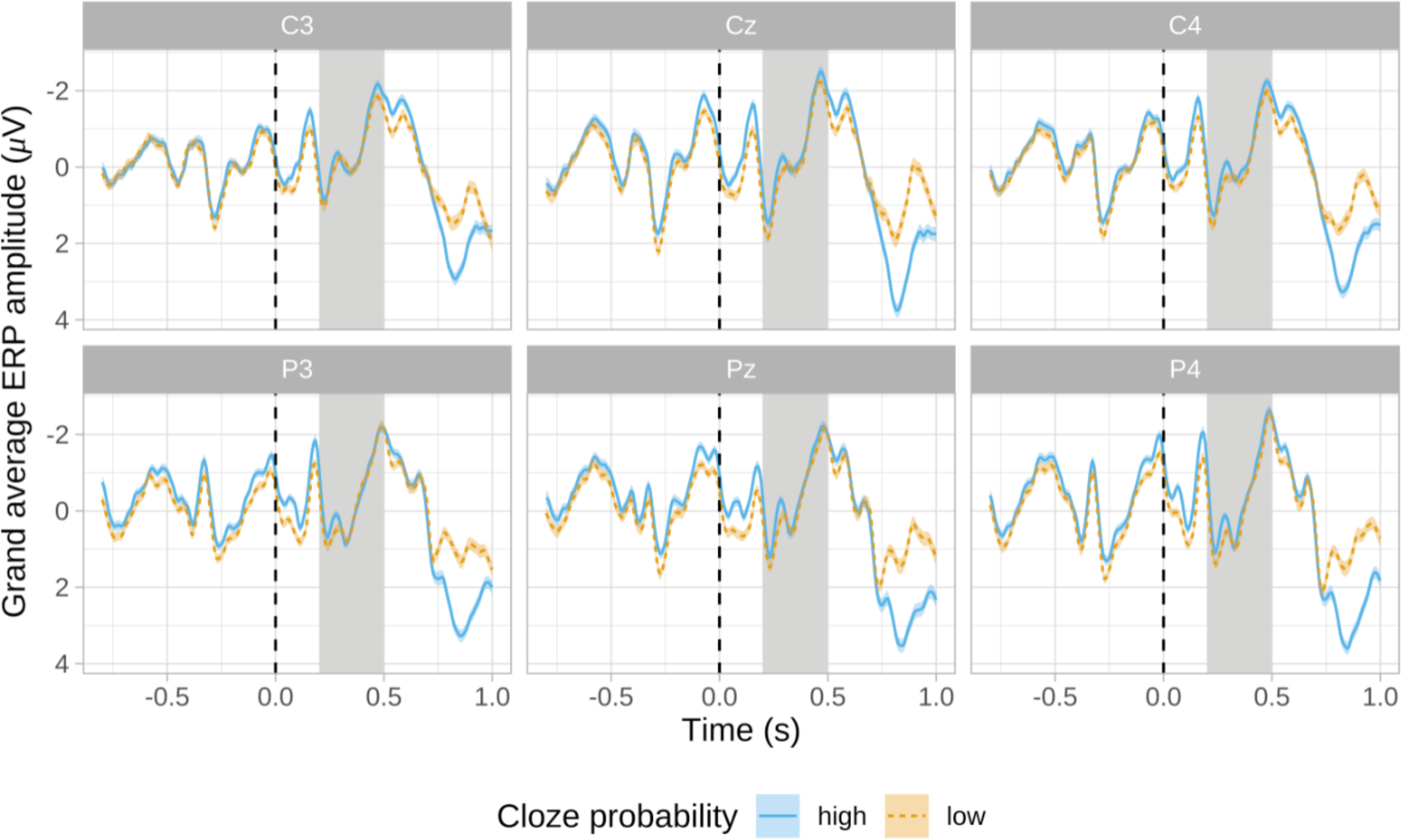
Grand average ERP plots of N400 effects for articles over channels of interest, according to article cloze probability (*n* = 312). Blue lines represent high article cloze probability (>50) whilst orange dashed lines reflect low article cloze probability (<50). Blue and orange shading depicts the standard errors for the two conditions. The N400 window is depicted via grey shading, and the black dashed line represents the onset of the article word. Note that the plot depicts non-baseline corrected ERP activity.

**Figure 6.**
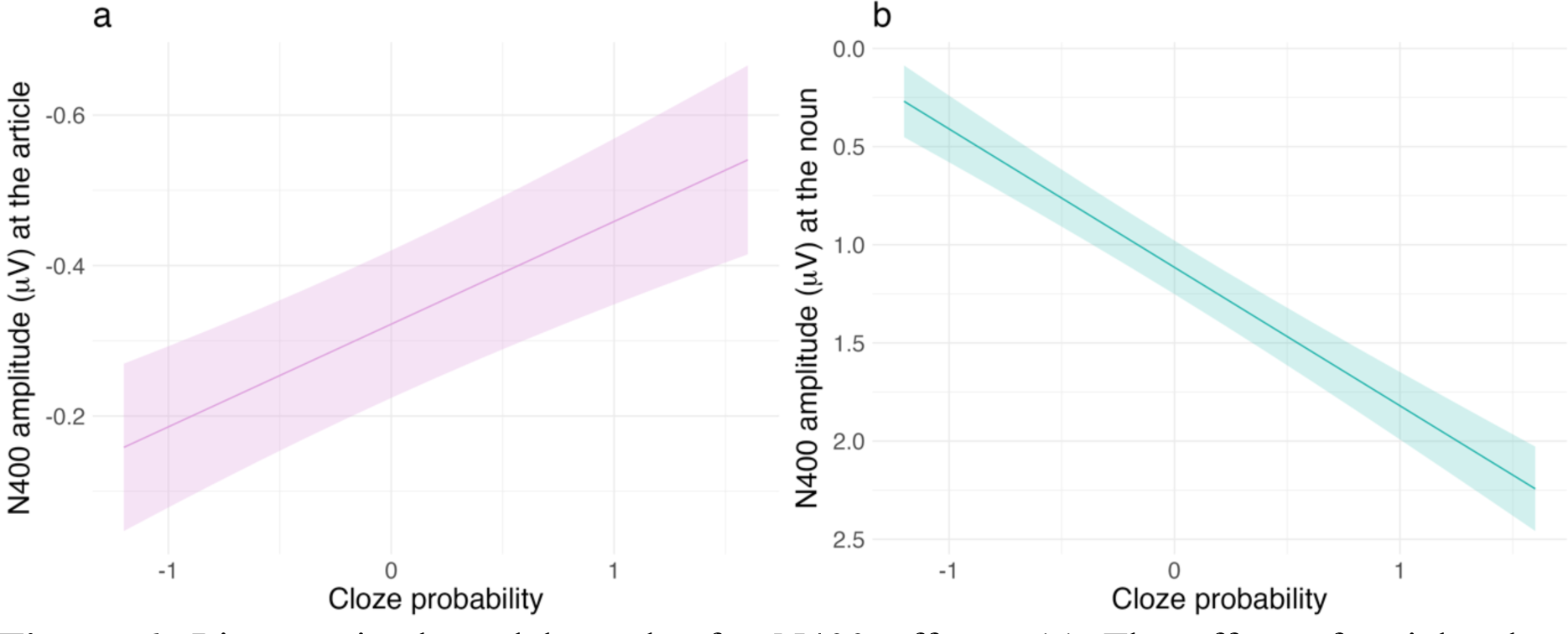
Linear mixed model results for N400 effects. **(a)** The effect of article cloze probability (scaled and centered) on N400 amplitude time-locked to articles (*p* <.001, β = - 0.14, *SE =* 0.03, 95% CI = -0.47 – -0.19). **(b)** The effect of noun cloze probability (scaled and centered) on N400 amplitude time-locked to nouns (*p* <.001, β = 0.71, *SE =* 0.07, 95% CI = 0.56 – 0.85). Shading in both plots reflects 83% confidence intervals and lines indicate model fit.

The effect of noun cloze probability, IAF and 1/f slope on the N400 amplitude following the nouns was explored in Model 4. The model detected a significant main effect of noun cloze probability on N400 amplitude time-locked to the noun (*p* <.001, β = 0.71, *SE =* 0.07, 95% CI = 0.56 – 0.85), where increases in noun cloze probability were associated with decreased (i.e., less negative) post-noun N400 amplitudes. The interaction between noun cloze and IAF on post-noun N400 amplitude did not reach significance (*p* = 0.395, β = -0.03, *SE =* 0.04, 95% CI = -0.11 – 0.04). The effect of noun cloze and 1/f slope was also nonsignificant (*p* = 0.127, β = 0.06, *SE =* 0.04, 95% CI = -0.02 – 0.13). Therefore, whilst cloze probability influenced N400 amplitudes following the articles and nouns, this relationship was not modulated by individual factors.

### Results of models with surprisal

As described above, whilst we ran SPP models with surprisal, the results either yielded the same outcomes as the cloze probability models or did not reveal interactions with the surprisal of the critical word. Therefore, for simplicity’s sake, the models are not reported here although the outputs are presented in *Appendix 3*.

Grand average N400 ERPs according to article surprisal are presented in *Figure 7,* whilst grand averages according to noun surprisal are shown in *Figure 8.* To determine whether article surprisal influenced the N400 amplitude directly following the article, Model 5 was computed, revealing a nonsignificant main effect of article surprisal on post-article N400 amplitude (*p* = 0.124, β = -0.08, *SE* = 0.05, 95% CI = -0.18 – 0.02). The interaction between article surprisal and IAF on N400 amplitude time-locked to the articles was nonsignificant (*p* = 0.557, β = - 0.02, *SE* = 0.04, 95% CI = -0.09 – 0.05). The effect of article surprisal and 1/f slope also did not reach significance (*p* = 0.671, β = -0.02, *SE* = 0.04, 95% CI = -0.09 – 0.06). Therefore, article surprisal, IAF and 1/f slope did not interact to influence the amplitude of the N400 component following the articles.

**Figure 7.**
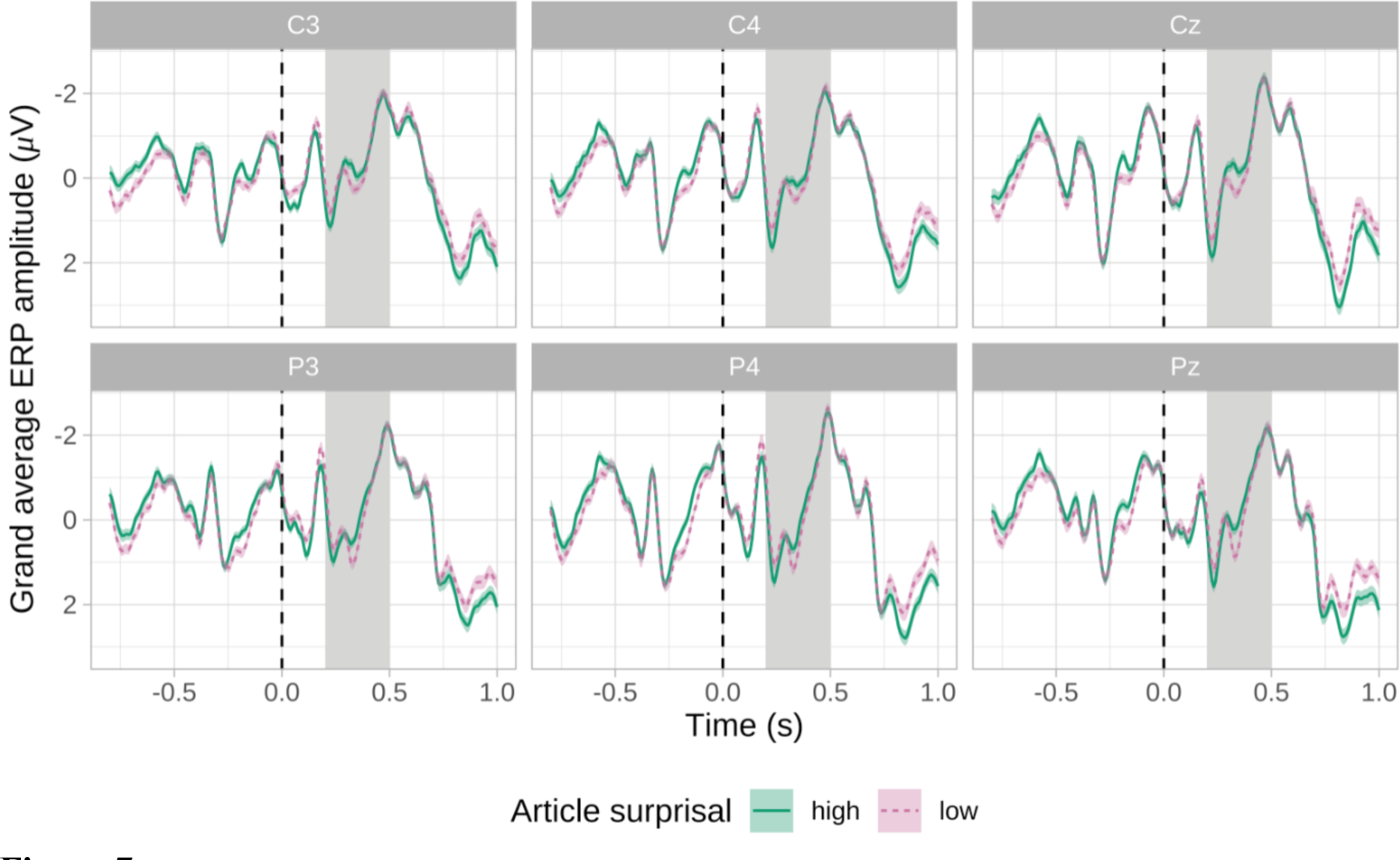
Grand average N400 ERP plots for articles over channels of interest, according to article surprisal (*n* = 312). Green lines represent high article surprisal (>4.08; the mean article surprisal value) whilst magenta dashed lines reflect low article surprisal (<4.08). Green and magenta shading depicts the standard errors for the two conditions. The N400 window is depicted via grey shading, and the black dashed line represents the onset of the article. Note that the plot depicts non-baseline corrected ERP activity.

**Figure 8.**
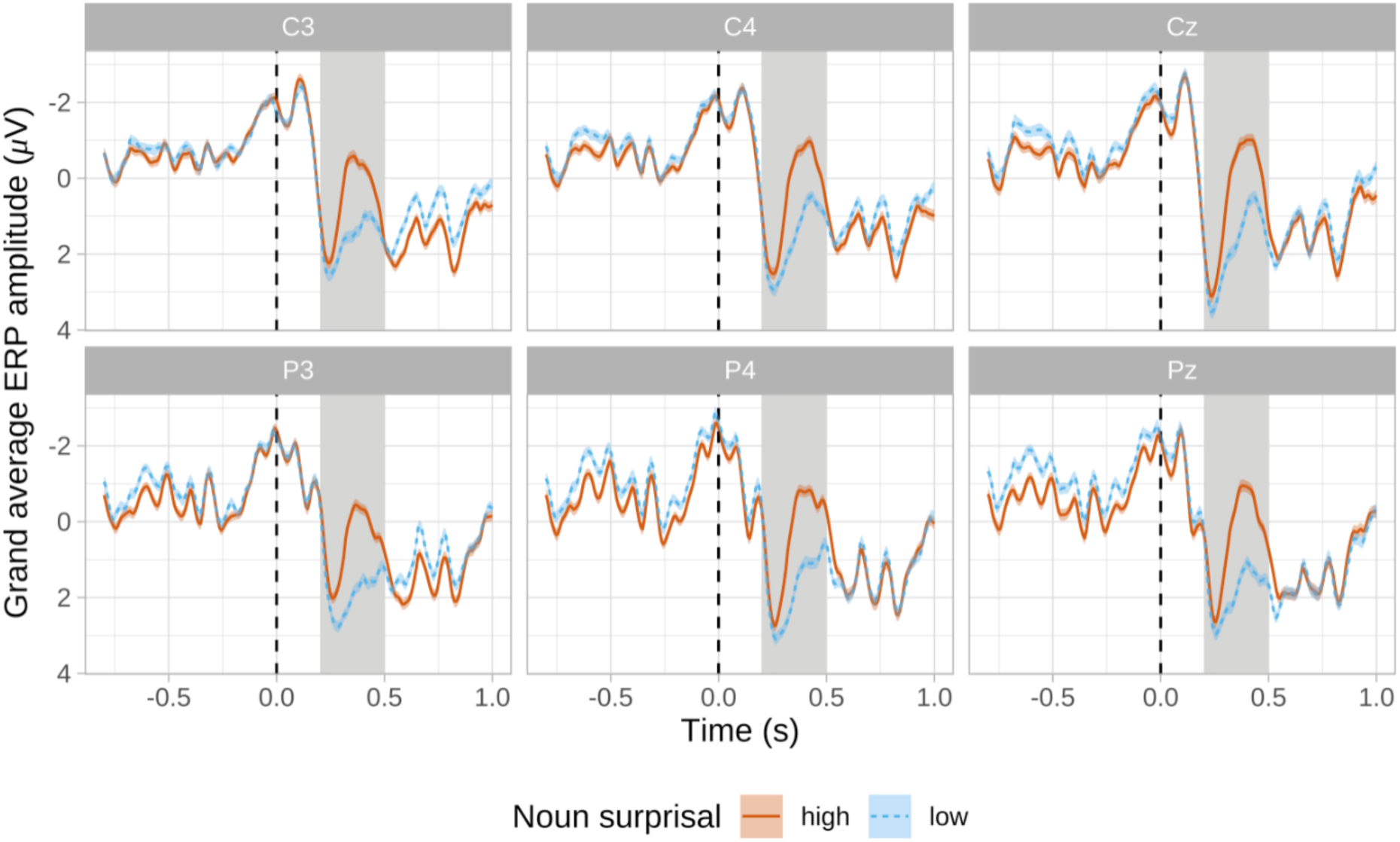
Grand average N400 ERP plots for nouns over channels of interest, according to noun surprisal (*n* = 312). Red lines represent high noun surprisal (>5.8; the mean noun surprisal value) whilst blue dashed lines reflect low noun surprisal (<5.8). Red and blue shading depicts the standard errors for the two conditions. The N400 window is depicted via grey shading, and the black dashed line represents the onset of the noun. Note that the plot depicts non-baseline corrected ERP activity.

To investigate the role of prior surprisal in prediction, Model 6a was fitted with article, a-1 and a-2 surprisal, and the 1/f slope. A significant interaction effect between article surprisal, a-1 surprisal and a-2 surprisal on post-article N400 amplitude was detected (*p* = 0.035, β = 0.11, *SE* = 0.05, 95% CI = 0.01 – 0.22). In most cases, increases in article surprisal were associated with increased (i.e., more negative) N400 amplitudes following the articles. However, this relationship was not observed when a-2 and a-1 surprisal were low (See *Figure 9a*). Model 6b predicted N400 amplitude from article, a-1 and a-2 surprisal, and IAF, which led to a significant interaction between article surprisal, a-2 surprisal and IAF on post-article N400 amplitude (*p* = 0.045, β = -0.07, *SE* = 0.04, 95% CI = -0.15 – -0.00). While increases in article surprisal were associated with larger (i.e., more negative) N400 amplitudes after the articles, this relationship was strongest for high IAF individuals when a-2 surprisal was high (See *Figure 9b*).

**Figure 9.**
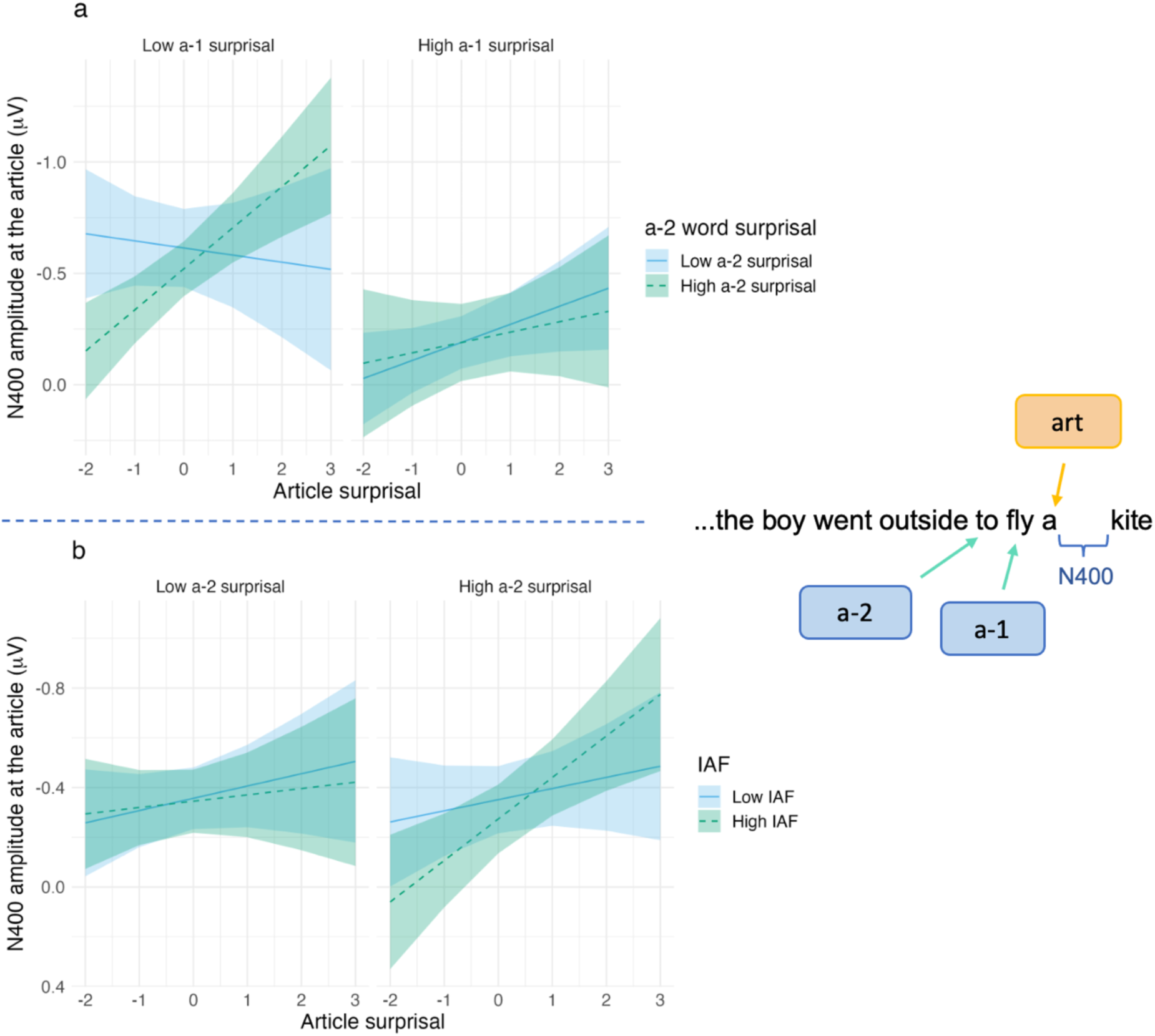
Linear mixed model results of article surprisal effects. **(a)** N400 amplitude following the articles, predicted by article–2 (a-2), article-1 (a-1), and article surprisal (*p* = 0.035, β = 0.11, *SE* = 0.05, 95% CI = 0.01 – 0.22). **(b)** N400 amplitude time-locked to articles, predicted by article-2 (a-2) surprisal, article surprisal and IAF (*p* = 0.045, β = -0.07, *SE* = 0.04, 95% CI = -0.15 – -0.00). Shading in both plots reflects 83% confidence intervals, lines indicate model fit, and the dichotomisation of variables is for visualisation purposes only. Note that both plots depict scaled and centered predictors.

The effect of noun surprisal on N400 amplitude time-locked to the nouns was investigated via Model 7. A significant main effect of noun surprisal on post-noun N400 amplitude was observed (*p* <.001, β = -0.79, *SE* = 0.12, 95% CI = -1.03 – -0.55), such that greater noun surprisal values were associated with larger (i.e., more negative) N400 amplitudes following the nouns.

### Correlational models

Shapiro-wilk tests revealed that article cloze probability and article surprisal were not normally distributed, and as such, a Spearman correlation was computed to examine the relationship between the two variables. The test revealed a significant but weak correlation (*r*(158) = -.25, *p* = 0.001), such that increases in article cloze were associated with decreases in article surprisal (see *Figure 10a*). A second Spearman correlation was calculated to determine the relationship between noun surprisal and noun cloze probability, after Shapiro-wilk tests revealed that both variables also did not conform to a normal distribution. Noun surprisal and noun cloze probability were significantly negatively correlated (*r*(157) = -.63, *p* <.001), with increases in noun cloze associated with decreases in noun surprisal (see *Figure 10b*). Finally, we examined the correlation between the 1/f slope and IAF using a Spearman correlation, as these variables were not normally distributed. The test revealed a significant but weak negative correlation between the two variables (*r*(310) = -.17, *p* <.01), such that increases in IAF were associated with decreased (flatter) 1/f slopes.

**Figure 10.**
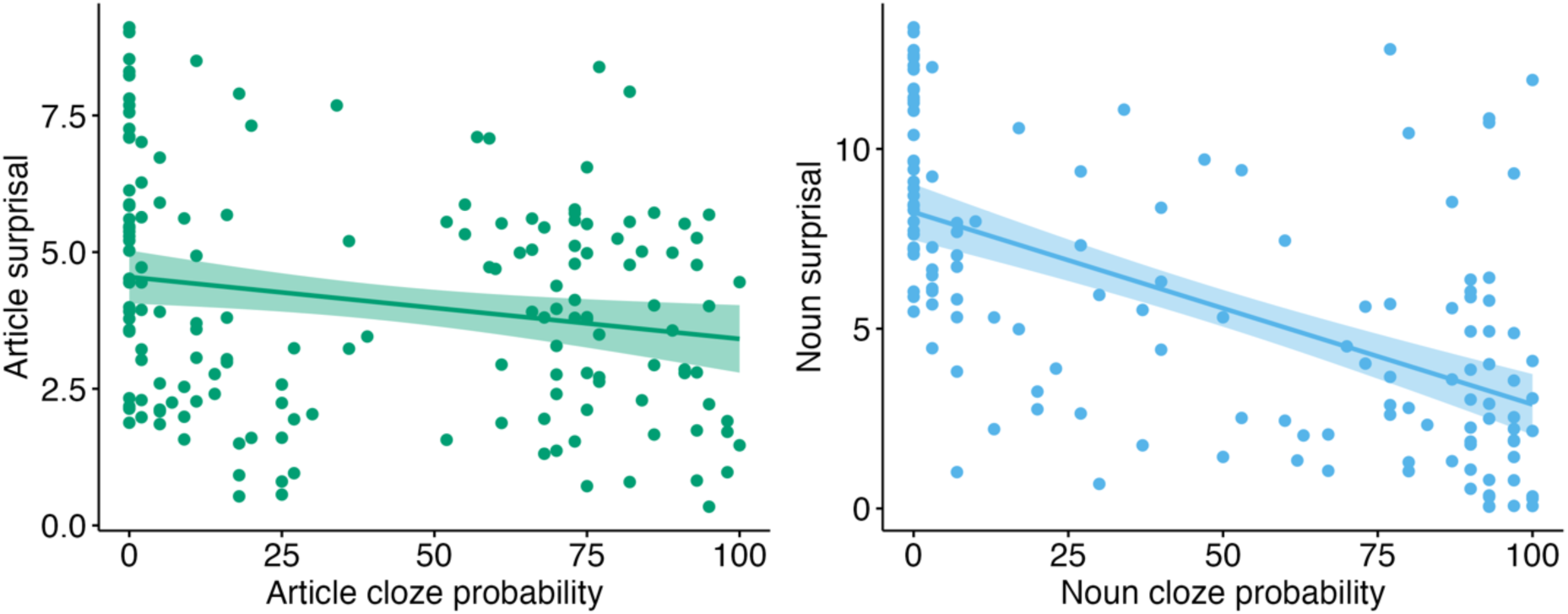
Spearman correlations depicting the relationship between cloze probability and surprisal. **(a)** The relationship between article cloze probability and article surprisal (*r*(158) = -.25, *p* = 0.001). **(b)** The relationship between noun cloze probability and noun surprisal (*r*(157) = -.63, *p* <.001).

## Discussion

The present study sought to investigate the degree to which prediction underlies linguistic processing via an analysis of individual neural differences and ERP activity in the Nieuwland et al. (2018) data. Our analysis did not reveal a positive relationship between word predictability and the magnitude of the SPP as was hypothesized. This may question the extent to which the SPP reflects prediction during natural language processing. Initial N400 models time-locked to articles demonstrated increases in N400 amplitude as article cloze probability increased, whilst article surprisal and post-article N400 amplitudes were not significantly related. At a first glance, this may be taken to suggest that the brain does not predict the phonological form of an upcoming noun, in line with Nieuwland and colleagues’ original findings. However, evidence for prediction arose when prior surprisal was considered, such that N400 amplitudes following the articles increased as article surprisal increased. This relationship was not observed when the surprisal of the two prior words was low but was strengthened for high IAF individuals when article-2 surprisal was high. Broadly, our reanalysis contributes novel insights to the debate surrounding prediction in language, by suggesting that prediction during linguistic comprehension depends on contextual and individual factors. This emphasises the adaptive and between-subject nature of prediction, whilst highlighting the variability that surrounds predictive processing at the article.

### The SPP may be a limited index of prediction during natural language processing

It was hypothesized that SPP amplitudes would increase (i.e., become more negative) as words became more predictable, reflecting anticipatory processing during linguistic comprehension. However, our analyses in the pre-stimulus window did not reveal a significant effect of article cloze probability on pre-article SPP amplitude, whilst SPP amplitudes preceding the nouns decreased (i.e., became more positive) as noun cloze probability increased. These findings are inconsistent with prior literature, which demonstrates a positive relationship between the magnitude of the SPP/SPN and word predictability (Grisoni et al., 2017, 2021; León-Cabrera et al., 2017, 2019). Whilst at face level, these findings might be taken to suggest that participants were not predicting upcoming semantic information, the results may be further illuminated by considering paradigm-specific effects.

As the SPP is proposed to reflect the prediction of semantic information (Grisoni et al., 2021; Pulvermüller & Grisoni, 2020), the observed null article finding may be explained by the overlap in meaning between the articles ‘a’ and ‘an,’ resulting in a similar semantic prediction between the two. Alternatively, the noun effect may suggest that SPP analyses were influenced by the temporal overlap between the pre-stimulus and N400 windows following the prior words. Most studies observing SPP/SPN effects employed intervals preceding the critical words of 1000ms or longer to capture activity independent of ERP components following the prior word (Grisoni et al., 2017, 2021; León-Cabrera et al., 2017, 2019). In contrast, the present study included interstimulus intervals of 300ms, which overlapped with the prior word’s N400 window. To test whether the N400 following the prior words was probed in our SPP analyses, we also ran the SPP models whilst controlling for prior word surprisal (for model outputs, see *Appendix 3*). This did not influence the results, likely because whilst the windows for the two components overlapped, the topographies differed (frontal for the SPP and centroparietal for the N400). Further, the relationship between critical word predictability (as measured by both cloze and surprisal) and SPP amplitude was not influenced by prior word surprisal or individual factors. This is in contrast with the N400 patterns at the article, which may provide evidence for probabilistic prediction under certain circumstances. Therefore, as an SPP effect in the expected direction was absent even after including factors that influenced prediction (as evidenced by the N400 effects), this may suggest that the SPP does not reflect naturalistic linguistic prediction. This raises the question of whether the longer intervals in other studies encourage prediction to a greater extent, or whether they give rise to a process that differs from natural language comprehension. Supporting this enquiry is a study by Kaan and Carlisle (2014), which presented participants with predictable and random letter strings and varied the length of the delay between the letters. The authors observed a negativity resembling the SPN/SPP only during longer delay trials, which was larger (more negative) for predictable versus random sequences (Kaan & Carlisle, 2014). This further supports the notion that such pre-stimulus negativities may manifest in the presence of longer delays, rendering the extent to which they mirror predictive processes unclear. Ultimately, while we sought to gauge prediction more directly via an analysis of the SPP, effects relating to the N400 may contribute more strongly to understandings of prediction in language.

### N400 effects at the article are variable and offer mixed results for prediction

Initial article analyses revealed that as article cloze probability increased, N400 amplitudes following the articles became larger (i.e., more negative), questioning the extent to which the brain predicts upcoming linguistic information. This result opposes both DeLong et al. (2005), who found the reversed effect, and Nieuwland et al. (2018), whose N400-article analysis did not reach significance (although the trend was detected in some laboratories in Nieuwland and colleagues’ replication analysis). Differences in the data analysis pipeline between our analysis and Nieuwland et al.’s (2018) study may have contributed to the differential results. Variations in pre-processing steps and our inclusion of pre-stimulus activity as a model covariate (which is shown to improve statistical power over baseline correction; Alday, 2019), may provide a potential explanation for this outcome. Interestingly, the results at the article differed depending on whether surprisal or cloze probability was used, highlighting the potential discrepancy between the two measures in their ability to gauge the predictability of function words such as articles.

When article surprisal was used to operationalize word predictability as opposed to article cloze probability, the N400 effect no longer reached significance. This is in line with Nieuwland and colleagues’ original analysis, suggesting that phonological prediction in language may be limited. On the other hand, it is inconsistent with Yan et al.’s (2017) reanalysis of Nieuwland et al. (2018), which revealed increases in the N400 as article surprisal increased (supporting phonological prediction in language). However, whilst we calculated surprisal using corpus-derived GPT-2 models, Yan and colleagues (2017) used the log-transformation of cloze as a measure of surprisal. Consequently, the mixed results at the article may in part arise from the use of differential word predictability metrics; a suggestion that is supported by the weak negative correlation between article cloze probability and article surprisal observed in the present analysis. Cloze probability and GPT-2 measures appear to model N400 patterns slightly differently, particularly at the lower end of the predictability spectrum (Szewczyk & Federmeier, 2022). Whilst cloze is limited in modelling N400 effects to unexpected words with cloze probabilities closer to zero, GPT-2 measures may explain more variance in N400 amplitudes here due to their reliance on the log scale (Szewczyk & Federmeier, 2022). Other research however suggests that cloze probability is a better predictor of N400 amplitude than GPT-2 surprisal (Michaelov et al., 2022). Interestingly, all noun analyses revealed a consistent effect in the typical direction (similarly to DeLong et al., 2005 and Nieuwland et al., 2018), and noun cloze probability and noun surprisal were strongly negatively related (in line with theory; Pickering & Gambi, 2018). This may also suggest that the predictability of function words such as articles is more difficult to gauge, potentially because such words are not as semantically rich as content words (e.g., nouns). Whilst semantic information accumulates across a sentence to constrain subsequent word choices, early speech sounds do not strongly limit later word possibilities, which may render predictability measurements difficult (Huettig et al., 2022). Therefore, whilst the present N400 results at a superficial level appear to provide evidence against phonological prediction, the diversity in outcomes at the article underscores the substantial variability that may surround the processing of this word. This is further emphasised by the observation that the relationship between article surprisal and post-article N400 amplitude appears subject to individual and contextual variation.

### Predictive processing depends on prior contextual information

While the above results may question the extent of prediction in language, the analyses including prior word surprisal suggest that predictive processing depends on prior context. When the surprisal of the two preceding words in the sentence was included in the statistical models, increases in article surprisal were associated with increases in N400 amplitude following the article (in line with DeLong et al., 2005). However, this effect was not present when the surprisal of the two prior words was low, suggesting that the N400 is sensitive to prior contextual information. This notion is broadly compatible with past research, which manipulates the previous context experimentally, consequently changing the predictability of the critical words (Boudewyn et al., 2015; Fleur et al., 2020; Freunberger & Roehm, 2017; Maess et al., 2016; Ness & Meltzer-Asscher, 2018; Szewczyk & Wodniecka, 2020; Szewczyk et al., 2022). Often, cloze probability at the critical words is used to gauge contextual effects (e.g., Maess et al., 2016; Federmeier et al., 2007). In contrast, the current analysis directly quantified the prior context in a continuous fashion, highlighting the importance of prior predictability as it unfolds naturally and incrementally over the course of a sentence. Whilst this analysis may not directly inform the underlying mechanisms behind these contextual effects, establishing that prior predictability is relevant for this relationship has value for the literature as it may form the basis of future, more in-depth explanations. Therefore, the observation of an N400 effect following the article when accounting for prior predictability may thus suggest that participants were predicting the phonological characteristics of the following noun, although only under certain circumstances. Nevertheless, it offers stronger support for the N400 as reflecting prediction versus integration. The shared semantic meaning of ’a/an’ in this context should not cause challenges in semantic integration and, consequently, should not result in an N400 effect akin to the one observed in the present study (DeLong et al., 2005). While this may thus support prediction perspectives of the N400, the absence of an article effect in cases of low prior word surprisal might be taken to suggest that participants were not predicting under such circumstances. At a first glance, this result appears somewhat counterintuitive: shouldn’t predictions theoretically be strongest in this case, given that prior context provides converging support for the current predictive model?

An alternative explanation for this effect may stem from the notion that the N400 reflects a precision-weighted prediction error signal (Bornkessel-Schlesewsky & Schlesewsky, 2019), with precision reflecting the amount of certainty (or inverse of variance) in the data (Feldman & Friston, 2010; Friston, 2005). According to Bayesian frameworks, precision influences the relative weightings of the sensory input vis-à-vis the prior belief, which has consequences for predictive model updating (Adams et al., 2013). In the present study, low prior word surprisal may have fostered highly precise predictions, thereby concomitantly decreasing the precision attributed to the article such that it was deemed irrelevant for hypothesis testing. This may have consequently resulted in weaker N400 amplitudes as prediction errors at the article were ‘ignored’ in favour of predictions. The potential for articles to be overlooked is compatible with prior eye-tracking findings, where shorter words are typically skipped more frequently than longer words (Slattery & Yates, 2018), with highly common two-letter words being skipped by skilled readers at a rate of approximately 75% during reading (Leinenger & Rayner, 2017; Rayner & McConkie, 1976 in Faber et al., 2020). Further, whilst word skipping is also influenced by predictability, such that highly predictable words are more likely to be skipped (e.g., Balota et al., 1985; Ehrlich & Rayner, 1981; Slattery & Yates, 2018), these effects appear to differ for articles. For example, Angele and Rayner (2013) presented participants with a preview of the article ‘the’ in parafoveal vision, in a manner that rendered it syntactically anomalous given the prior sentence context. Participants tended to skip the preview despite its poor fit with the sentence, suggesting that the anomalous article was overlooked. Taken together, these findings may suggest that articles are frequently skipped, even when unpredictable. Furthermore, readers with greater print exposure and spelling ability have been shown to skip words more frequently (Faber et al., 2020; Slattery & Yates, 2018), highlighting the role of individual variability in this relationship. Therefore, even though results from natural reading are, of course, not directly comparable to the word-by-word presentation employed in the present experiment, it appears possible that article-induced prediction errors were not considered here under conditions of high prior predictability, analogous to the article skipping findings. In addition to providing a potential explanation for the reduced N400 amplitudes that we observed, this notion is compatible with active inference theories, which propose that prediction error is not only reduced via changes to one’s predictive model, but through active sampling of the sensory environment in accordance with one’s hypothesis (Friston et al., 2016; Friston & Frith, 2015; Parr et al., 2022). As such, comprehenders may at times neglect unexpected articles in favour of a top-down expectation.

Ultimately, the present results, in addition to the mixed findings regarding cloze and surprisal described above, point towards a uniqueness within the article that gives rise to significant variability, and which may question the utility of this and similar paradigms. If information at the article is overlooked under certain circumstances, and if the relative weighting of the bottom-up versus top-down information influences the N400, interpretations from such paradigms may be more complicated than initially assumed. Interestingly, this could also suggest that the original appeal of Delong et al.’s (2005) design – that the article does not contain rich lexical-semantic content of its own – may simultaneously be its greatest weakness. Further, the results of the present study align with the proposal that pre-nominal prediction effects may be small and difficult to detect (Kochari & Flecken, 2019; Nicenboim et al., 2020), as a pattern in the expected direction was only observed when prior predictability was considered. This suggests that the difficulty in observing the trend could arise from the inherent variability surrounding processing at the article (as well as at similar pre-nominal words). In further support of the variability surrounding the article is the observation that this relationship was influenced by IAF.

### Individual alpha frequency (IAF) modulates predictive processing during language comprehension

In addition to effects of prior context, N400 amplitude following the article was influenced by IAF, emphasising the subject-specific nature of prediction. Whilst the post-article N400 amplitude increased (i.e., became more negative) with increasing article surprisal, this trend was strongest for high IAF individuals when article-2 surprisal was high. Therefore, it is possible to suggest that under conditions of uncertainty, high IAF individuals rely on prediction to a greater extent than low IAF individuals. Alternatively, and in line with our previous discussions concerning precision, this result may also imply that high IAF individuals were weighting the bottom-up information at the article more strongly (i.e., they were affording more precision to the article), resulting in stronger post-article N400 effects. As high prior surprisal likely renders it difficult to form a precise prediction, such individuals may have prioritised the sensory input for model updating. This is supported by prior research, where those with a high IAF demonstrate faster information processing speeds, suggesting that they may sample the perceptual environment more rapidly (Klimesch et al., 1996; Samaha & Postle, 2015; Surwillo, 1961, 1963). Other research proposes that such individuals have a greater inclination towards predictive model updating (Kurthen et al., 2020), which may manifest in larger N400 amplitudes following surprising articles as observed here. Therefore, whilst N400 effects time-locked to the article are typically discussed as reflecting prediction for the upcoming noun, the components’ ties to precision (Bornkessel-Schlesewsky & Schlesewsky, 2019) imply that high IAF individuals may adopt a more flexible model updating strategy during conditions of uncertainty. However, further research would be beneficial to determine whether IAF directly influences stimulus precision weightings, potentially by investigating correlates that index prediction errors for precision weightings themselves (e.g., the P300 ERP component, as suggested by Bornkessel-Schlesewsky, Alday, et al., 2022). Additionally, whilst IAF influenced N400 amplitudes, we did not detect any effects relating to the 1/f aperiodic slope. Although we investigated 1/f as a between-subjects (trait-like) measure, in line with prior research (e.g., Bornkessel-Schlesewsky et al., 2022; Demuru & Fraschini, 2020; Dziego et al., 2023; McSweeney et al., 2021; Voytek & Knight, 2015), the 1/f slope also shows state-like changes and has been linked with norepinephrine activity and phasic changes in neural gain (Pertermann et al., 2019; Rosenblum, 2023). Consequently, future research may benefit from examining 1/f on a trial-by-trial basis to determine whether changes relating to prediction may be more transient. Nevertheless, the present findings emphasise the role of IAF in predictive processing and suggest that whilst prediction may be an inherent property of cognition (Friston, 2010), the extent to which information is used for predictive model updating is subject to inter-individual variability.

## Conclusion

The present study sheds light on the adaptability of predictive processing, whilst emphasising the susceptibility of the N400 component to judgements concerning precision. Whilst we observed evidence for phonological linguistic prediction under certain circumstances, our results speak to the potential sources of variability that alter prediction tendencies at the article, implying that under conditions of high prior predictability, prediction errors at the article may be overlooked in favour of one’s predictive model. This may explain how internal language models cope with change as an inherent property of language, as prior context may inform the extent to which subsequent information is used for predictive model updating. The role of IAF in this relationship also suggests that some individuals may be more likely to adjust their predictions than others, having implications for understandings of prediction as a fundamental cognitive mechanism.

## Data availability statement

Nieuwland and colleagues’ data used in the present analysis is publicly available at https://osf.io/eyzaq/, whilst our data analysis scripts to replicate this investigation are publicly available at https://osf.io/b7txn/.

## Appendix 1: Model comparison tables

**Appendix 1 – Table 1.**
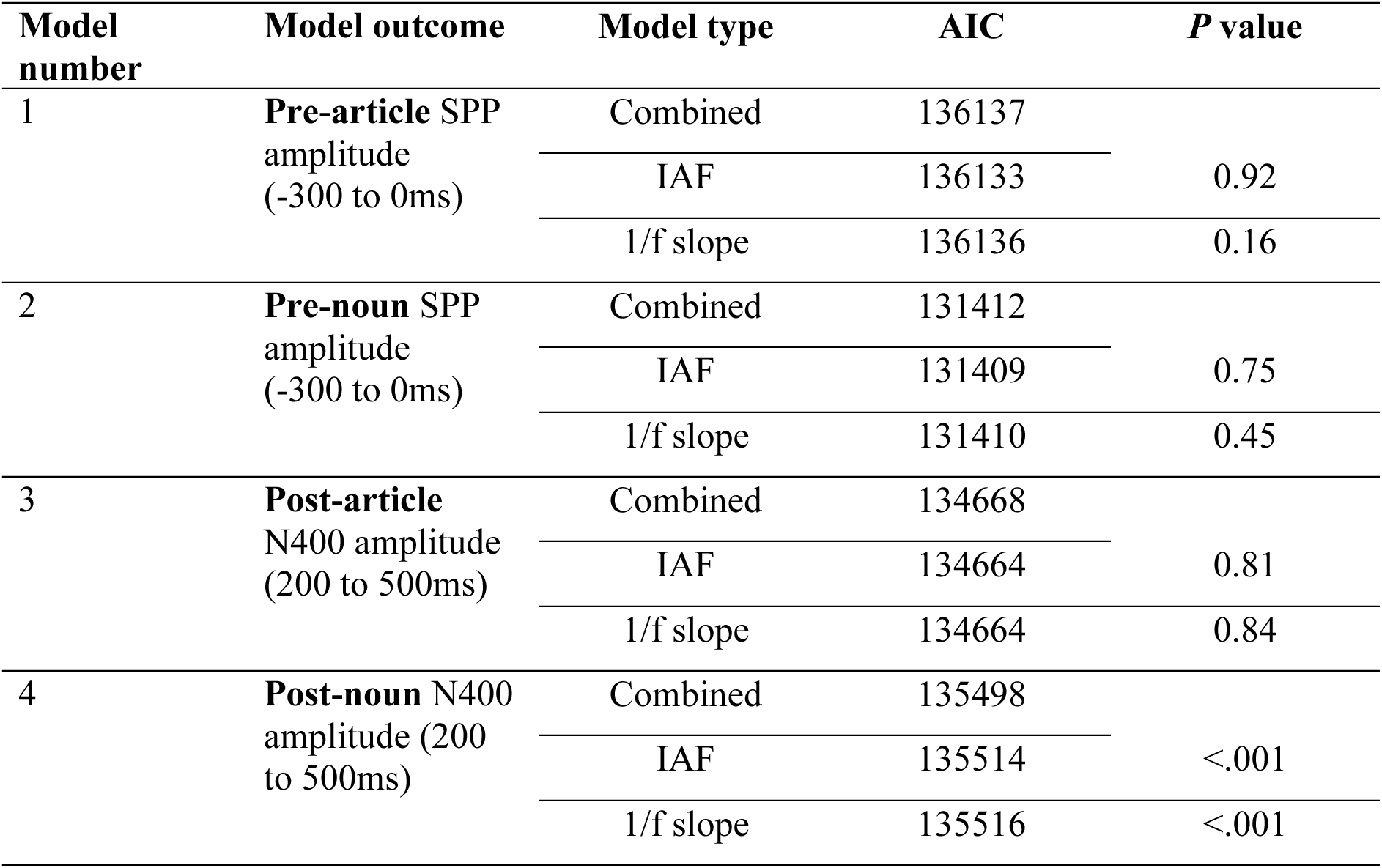
Akaike criterion (AIC) values for combined IAF and 1/f slope models, compared to separate IAF and 1/f slope models. P values reflect the comparison of the combined models’ AIC and the IAF and 1/f slope models’ AIC values, respectively.

**Appendix 1 – Table 2.** Results of likelihood ratio tests comparing basic model versus complex model including random slopes.

| Model number | Model type |  | P-value |
| --- | --- | --- | --- |
|  | Basic | Complex (with random slopes) |  |
| 2 | -65719 | -65707 | <.001 |
| 3 | -67340 | -67339 | 0.62 |
| 4 | -67834 | -67749 | <.001 |
| 5 | -67346 | -67336 | <.001 |
| 6a | -65695 | -65688 | <.01 |
| 6b | -65696 | -65688 | <.01 |
| 7 | -59292 | -59244 | <.001 |

## Appendix 2: Outputs of statistical models

**Appendix 2 – Table 1.** Model 1: SPP_amp_mean ∼ cloze_scale * iaf_scale + cloze_scale * slope_scale + baseline_amp_scale + (1|subjnr) + (1|item_no), data= art_locs_sum

| *Predictors* | **SPP_amp_mean** |  |  |  |
| --- | --- | --- | --- | --- |
|  | *Estimates* | *std. Error* | *CI (95%)* | *p* |
| (Intercept) | -0.00 | 0.12 | -0.23 – 0.23 | 0.993 |
| cloze scale | 0.06 | 0.13 | -0.18 – 0.31 | 0.620 |
| iaf scale | 0.12 | 0.08 | -0.04 – 0.28 | 0.157 |
| slope scale | 0.03 | 0.08 | -0.13 – 0.19 | 0.696 |
| baseline amp scale | -1.20 | 0.04 | -1.27 – -1.12 | **<0.001** |
| cloze scale × iaf scale | -0.03 | 0.08 | -0.19 – 0.13 | 0.748 |
| cloze scale × slope scale | -0.03 | 0.08 | -0.19 – 0.13 | 0.715 |
| **Random Effects** |  |  |  |  |
| $\sigma^2$ | 28.78 | | | |
| $\tau_{00}$ subjnr | 0.33 | | | |
| $\tau_{00}$ item_no | 0.16 | | | |
| ICC | 0.02 |  |  |  |
| N<sub>subjnr</sub> | 312 |  |  |  |
| N<sub>item_no</sub> | 80 |  |  |  |
| Observations | 21922 |  |  |  |
| Marginal R<sup>2</sup> / Conditional R<sup>2</sup> | 0.046 / 0.062 |  |  |  |

**Appendix 2 – Table 2.** Model 2: SPP_amp_mean ∼ cloze_scale * iaf_scale + cloze_scale * slope_scale + baseline_amp_scale + (1+cloze_scale|subjnr) + (1+cloze_scale|item_no), data= noun_locs_sum

| *Predictors* | **SPP_amp_mean** |  |  |  |
| --- | --- | --- | --- | --- |
|  | *Estimates* | *std. Error* | *CI (95%)* | *p* |
| (Intercept) | -1.04 | 0.07 | -1.18 – -0.90 | **<0.001** |
| cloze scale | 0.11 | 0.05 | 0.01 – 0.21 | **0.038** |
| iaf scale | -0.01 | 0.05 | -0.11 – 0.10 | 0.882 |
| slope scale | 0.04 | 0.05 | -0.06 – 0.14 | 0.449 |
| baseline amp scale | -0.54 | 0.04 | -0.61 – -0.46 | **<0.001** |
| cloze scale × iaf scale | -0.05 | 0.04 | -0.12 – 0.03 | 0.220 |
| cloze scale × slope scale | -0.00 | 0.04 | -0.08 – 0.07 | 0.919 |
| **Random Effects** |  |  |  |  |
| $\sigma^2$ | 27.58 | | | |
| $\tau_{00}$ subjnr | 0.43 | | | |
| $\tau_{00}$ item_no | 0.17 | | | |
| $\tau_{11}$ subjnr.cloze_scale | 0.07 | | | |
| $\tau_{11}$ item_no.cloze_scale | 0.08 | | | |
| $\rho_{01}$ subjnr | -0.20 | | | |
| $\rho_{01}$ item_no | -0.16 | | | |
| ICC | 0.03 |  |  |  |
| N subjnr | 312 |  |  |  |
| N item_no | 80 |  |  |  |
| Observations | 21282 |  |  |  |
| Marginal R<sup>2</sup> / Conditional R<sup>2</sup> | 0.011 / 0.037 |  |  |  |

**Appendix 2 – Table 3.** Model 3: N400_amp_mean ∼ cloze_scale * iaf_scale + cloze_scale * slope_scale + prestim_scale + (1|subjnr) + (1|item_no), data= combo_art_sum

| *Predictors* | N400_amp_mean |  |  |  |
| --- | --- | --- | --- | --- |
|  | *Estimates* | *std. Error* | *CI (95%)* | *p* |
| (Intercept) | -0.33 | 0.07 | -0.47 – -0.19 | **<0.001** |
| cloze scale | -0.14 | 0.03 | -0.20 – -0.07 | **<0.001** |
| iaf scale | 0.03 | 0.04 | -0.06 – 0.11 | 0.561 |
| slope scale | -0.00 | 0.04 | -0.08 – 0.08 | 0.985 |
| prestim scale | -1.71 | 0.03 | -1.77 – -1.65 | **<0.001** |
| cloze scale × iaf scale | 0.00 | 0.03 | -0.06 – 0.07 | 0.937 |
| cloze scale × slope scale | -0.02 | 0.03 | -0.09 – 0.04 | 0.512 |
| **Random Effects** |  |  |  |  |
| $\sigma^2$ | 21.95 | | | |
| $\tau_{00}$ subjnr | 0.24 | | | |
| $\tau_{00}$ item_no | 0.27 | | | |
| ICC | 0.02 |  |  |  |
| N<sub>subjnr</sub> | 312 |  |  |  |
| N<sub>item_no</sub> | 80 |  |  |  |
| Observations | 22669 |  |  |  |
| Marginal R<sup>2</sup> / Conditional R<sup>2</sup> | 0.115 / 0.135 |  |  |  |

**Appendix 2 – Table 4.** Model 4: N400_amp_mean ∼ cloze_scale * iaf_scale + cloze_scale * slope_scale + prestim_scale + (1+cloze_scale|subjnr) + (1+cloze_scale|item_no), data= combo_noun_sum

| *Predictors* | **N400_amp_mean** |  |  |  |
| --- | --- | --- | --- | --- |
|  | *Estimates* | *std. Error* | *CI (95%)* | *p* |
| (Intercept) | 1.11 | 0.10 | 0.92 – 1.31 | **<0.001** |
| cloze scale | 0.71 | 0.07 | 0.56 – 0.85 | **<0.001** |
| iaf scale | 0.15 | 0.07 | 0.01 – 0.29 | **0.039** |
| slope scale | -0.02 | 0.07 | -0.16 – 0.12 | 0.790 |
| prestim scale | -1.97 | 0.03 | -2.03 – -1.90 | **<0.001** |
| cloze scale × iaf scale | -0.03 | 0.04 | -0.11 – 0.04 | 0.395 |
| cloze scale × slope scale | 0.06 | 0.04 | -0.02 – 0.13 | 0.127 |
| **Random Effects** |  |  |  |  |
| $\sigma^2$ | 23.47 | | | |
| $\tau_{00}$ subjnr | 1.20 | | | |
| $\tau_{00}$ item_no | 0.38 | | | |
| $\tau_{11}$ subjnr.cloze_scale | 0.13 | | | |
| $\tau_{11}$ item_no.cloze_scale | 0.30 | | | |
| $\rho_{01}$ subjnr | 0.00 | | | |
| $\rho_{01}$ item_no | 0.04 | | | |
| ICC | 0.08 |  |  |  |
| N subjnr | 312 |  |  |  |
| N item_no | 80 |  |  |  |
| Observations | 22463 |  |  |  |
| Marginal R<sup>2</sup> / Conditional R<sup>2</sup> | 0.148 / 0.217 |  |  |  |

**Appendix 2 – Table 5.** Model 5: N400_amp_mean ∼ art_s_scale * iaf_scale + art_s_scale * slope_scale + prestim_scale + (1+art_s_scale|subjnr) + (1+art_s_scale|item_no), data=combo_art_sum

| *Predictors* | **N400_amp_mean** |  |  |  |
| --- | --- | --- | --- | --- |
|  | *Estimates* | *std. Error* | *CI (95%)* | *p* |
| (Intercept) | -0.33 | 0.07 | -0.47 – -0.19 | **<0.001** |
| art s scale | -0.08 | 0.05 | -0.18 – 0.02 | 0.124 |
| iaf scale | 0.02 | 0.04 | -0.06 – 0.11 | 0.564 |
| slope scale | 0.00 | 0.04 | -0.08 – 0.08 | 0.995 |
| prestim scale | -1.71 | 0.03 | -1.77 – -1.65 | **<0.001** |
| art s scale × iaf scale | -0.02 | 0.04 | -0.09 – 0.05 | 0.557 |
| art s scale × slope scale | -0.02 | 0.04 | -0.09 – 0.06 | 0.671 |
| **Random Effects** |  |  |  |  |
| $\sigma^2$ | 21.84 | | | |
| $\tau_{00}$ subjnr | 0.25 | | | |
| $\tau_{00}$ item_no | 0.25 | | | |
| $\tau_{11}$ subjnr.art_s_scale | 0.10 | | | |
| $\tau_{11}$ item_no.art_s_scale | 0.05 | | | |
| $\rho_{01}$ subjnr | -0.13 | | | |
| $\rho_{01}$ item_no | 0.27 | | | |
| ICC | 0.03 |  |  |  |
| N subjnr | 312 |  |  |  |
| N item_no | 80 |  |  |  |
| Observations | 22669 |  |  |  |
| Marginal R<sup>2</sup> / Conditional R<sup>2</sup> | 0.114 / 0.140 |  |  |  |

**Appendix 2 – Table 6.**
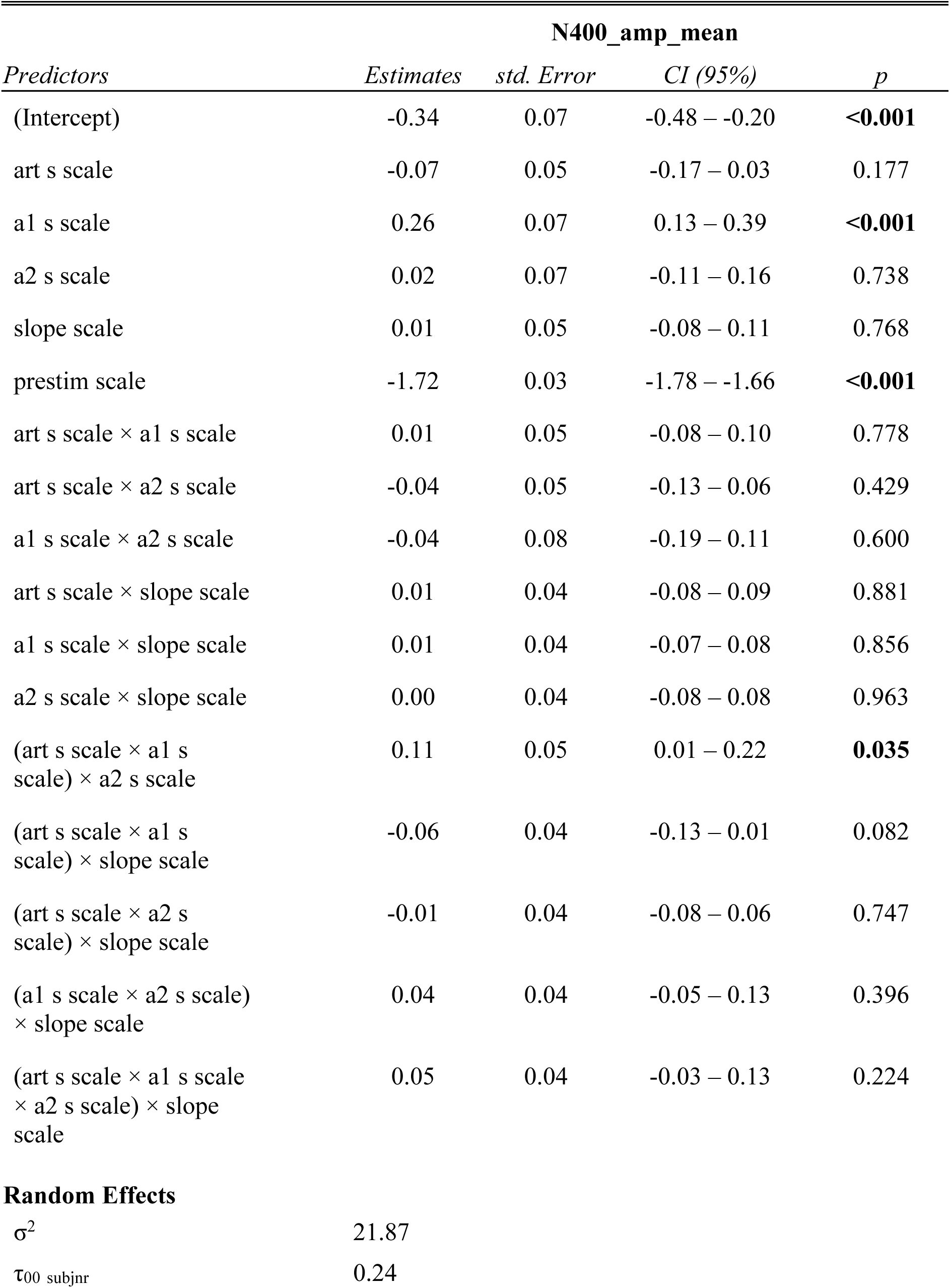
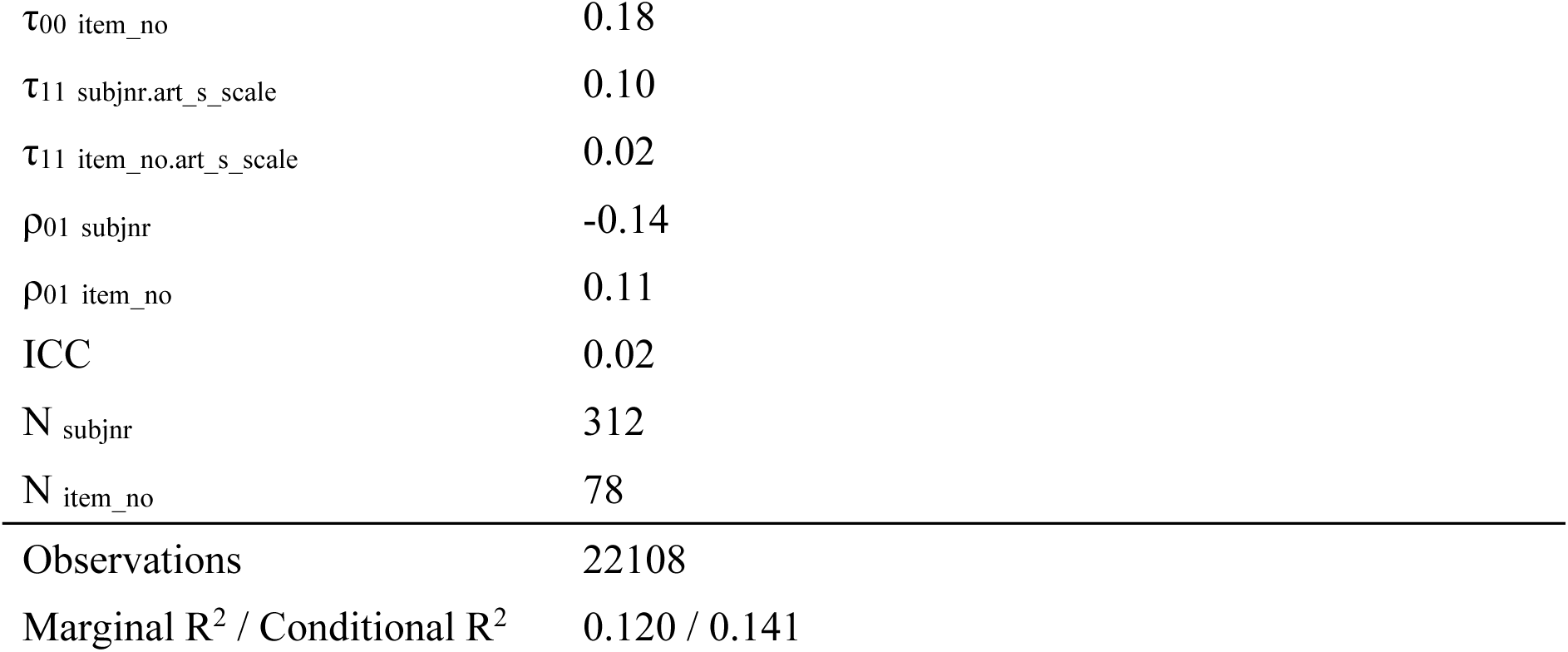
Model 6a: N400_amp_mean ∼ art_s_scale * a1_s_scale * a2_s_scale * slope_scale + prestim_scale + (1+art_s_scale|subjnr) + (1+art_s_scale|item_no), data=combo_art_sum

| *Predictors* | N400_amp_mean |  |  |  |
| --- | --- | --- | --- | --- |
|  | *Estimates* | *std. Error* | *CI (95%)* | *p* |
| (Intercept) | -0.34 | 0.07 | -0.48 – -0.20 | **<0.001** |
| art s scale | -0.07 | 0.05 | -0.17 – 0.03 | 0.177 |
| a1 s scale | 0.26 | 0.07 | 0.13 – 0.39 | **<0.001** |
| a2 s scale | 0.02 | 0.07 | -0.11 – 0.16 | 0.738 |
| slope scale | 0.01 | 0.05 | -0.08 – 0.11 | 0.768 |
| prestim scale | -1.72 | 0.03 | -1.78 – -1.66 | **<0.001** |
| art s scale × a1 s scale | 0.01 | 0.05 | -0.08 – 0.10 | 0.778 |
| art s scale × a2 s scale | -0.04 | 0.05 | -0.13 – 0.06 | 0.429 |
| a1 s scale × a2 s scale | -0.04 | 0.08 | -0.19 – 0.11 | 0.600 |
| art s scale × slope scale | 0.01 | 0.04 | -0.08 – 0.09 | 0.881 |
| a1 s scale × slope scale | 0.01 | 0.04 | -0.07 – 0.08 | 0.856 |
| a2 s scale × slope scale | 0.00 | 0.04 | -0.08 – 0.08 | 0.963 |
| (art s scale × a1 s scale) × a2 s scale | 0.11 | 0.05 | 0.01 – 0.22 | **0.035** |
| (art s scale × a1 s scale) × slope scale | -0.06 | 0.04 | -0.13 – 0.01 | 0.082 |
| (art s scale × a2 s scale) × slope scale | -0.01 | 0.04 | -0.08 – 0.06 | 0.747 |
| (a1 s scale × a2 s scale) × slope scale | 0.04 | 0.04 | -0.05 – 0.13 | 0.396 |
| (art s scale × a1 s scale × a2 s scale) × slope scale | 0.05 | 0.04 | -0.03 – 0.13 | 0.224 |
| **Random Effects** |  |  |  |  |
| $\sigma^2$ | 21.87 | | | |
| $\tau_{00}$ subjnr | 0.24 | | | |

|  |  |
| --- | --- |
| $\tau_{00}$ item_no | 0.18 |
| $\tau_{11}$ subjnr.art_s_scale | 0.10 |
| $\tau_{11}$ item_no.art_s_scale | 0.02 |
| $\rho_{01}$ subjnr | -0.14 |
| $\rho_{01}$ item_no | 0.11 |
| ICC | 0.02 |
| $N_{\text{subjnr}}$ | 312 |
| $N_{\text{item\_no}}$ | 78 |
| Observations | 22108 |
| Marginal $R^2$ / Conditional $R^2$ | 0.120 / 0.141 |

**Appendix 2 – Table 7.**
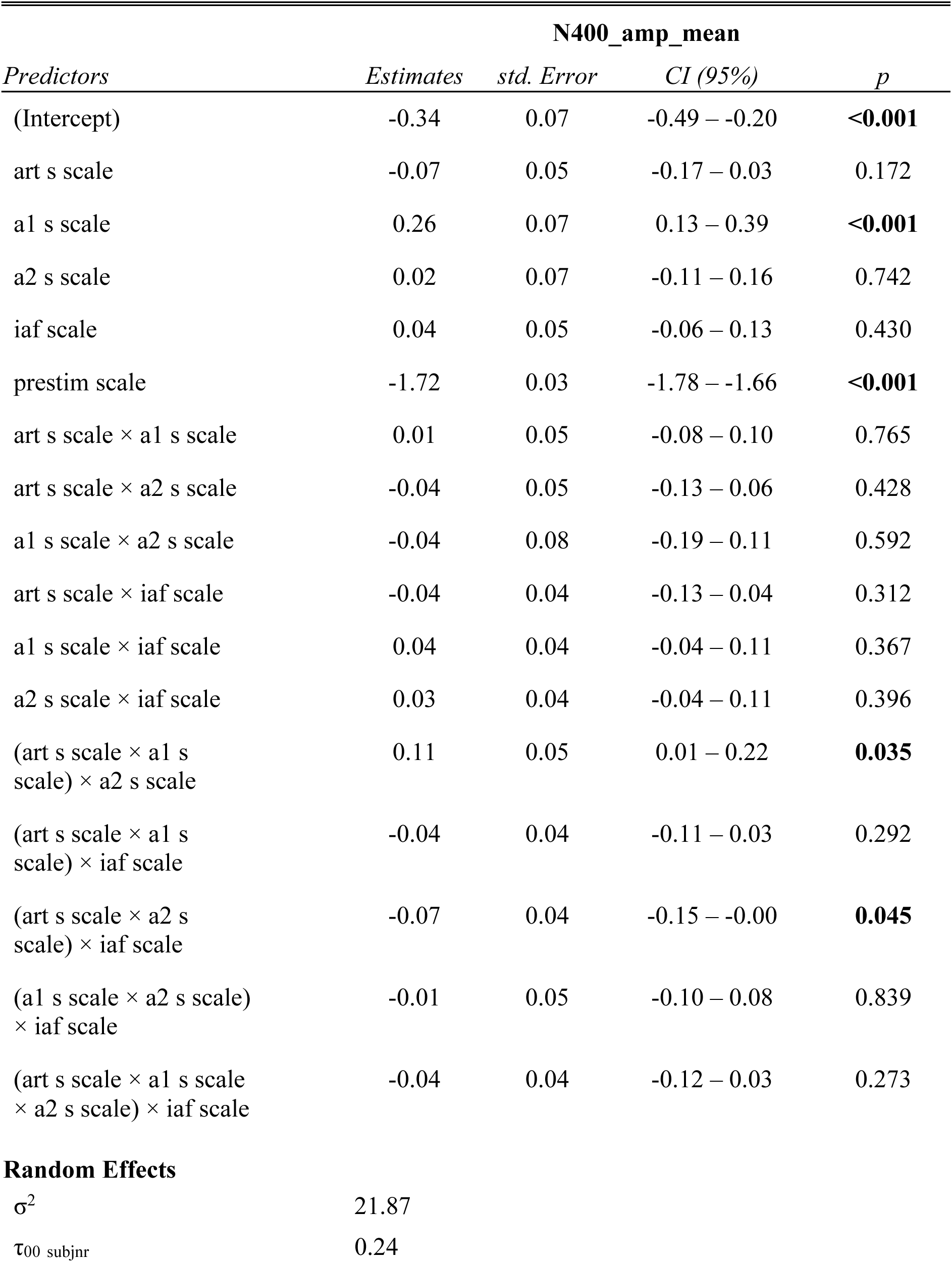
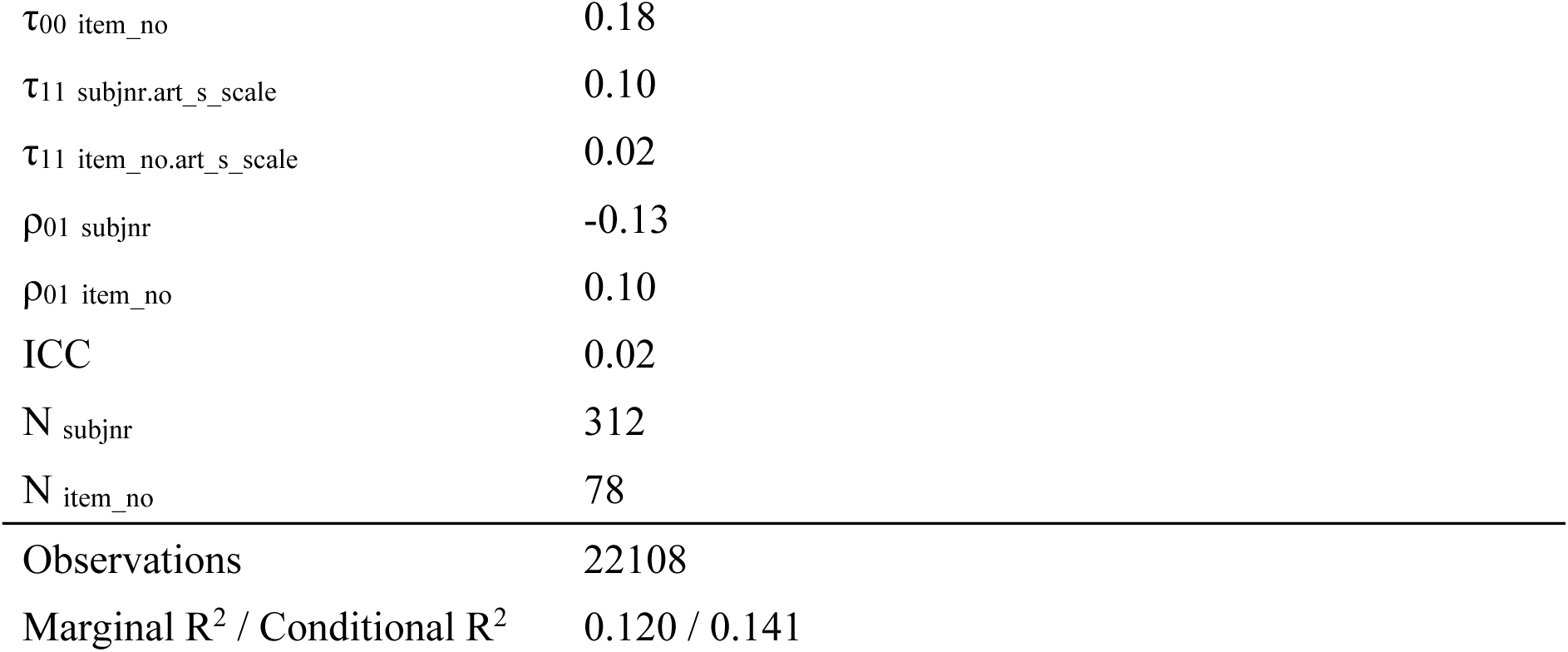
Model 6b: N400_amp_mean ∼ art_s_scale * a1_s_scale * a2_s_scale * iaf_scale + prestim_scale+ (1+art_s_scale|subjnr) + (1+art_s_scale|item_no), data=combo_art_sum

| *Predictors* | **N400_amp_mean** |  |  |  |
| --- | --- | --- | --- | --- |
|  | *Estimates* | *std. Error* | *CI (95%)* | *p* |
| (Intercept) | -0.34 | 0.07 | -0.49 – -0.20 | **<0.001** |
| art s scale | -0.07 | 0.05 | -0.17 – 0.03 | 0.172 |
| a1 s scale | 0.26 | 0.07 | 0.13 – 0.39 | **<0.001** |
| a2 s scale | 0.02 | 0.07 | -0.11 – 0.16 | 0.742 |
| iaf scale | 0.04 | 0.05 | -0.06 – 0.13 | 0.430 |
| prestim scale | -1.72 | 0.03 | -1.78 – -1.66 | **<0.001** |
| art s scale × a1 s scale | 0.01 | 0.05 | -0.08 – 0.10 | 0.765 |
| art s scale × a2 s scale | -0.04 | 0.05 | -0.13 – 0.06 | 0.428 |
| a1 s scale × a2 s scale | -0.04 | 0.08 | -0.19 – 0.11 | 0.592 |
| art s scale × iaf scale | -0.04 | 0.04 | -0.13 – 0.04 | 0.312 |
| a1 s scale × iaf scale | 0.04 | 0.04 | -0.04 – 0.11 | 0.367 |
| a2 s scale × iaf scale | 0.03 | 0.04 | -0.04 – 0.11 | 0.396 |
| (art s scale × a1 s scale) × a2 s scale | 0.11 | 0.05 | 0.01 – 0.22 | **0.035** |
| (art s scale × a1 s scale) × iaf scale | -0.04 | 0.04 | -0.11 – 0.03 | 0.292 |
| (art s scale × a2 s scale) × iaf scale | -0.07 | 0.04 | -0.15 – -0.00 | **0.045** |
| (a1 s scale × a2 s scale) × iaf scale | -0.01 | 0.05 | -0.10 – 0.08 | 0.839 |
| (art s scale × a1 s scale × a2 s scale) × iaf scale | -0.04 | 0.04 | -0.12 – 0.03 | 0.273 |
| **Random Effects** |  |  |  |  |
| $\sigma^2$ | 21.87 | | | |
| $\tau_{00}$ subjnr | 0.24 | | | |

|  |  |
| --- | --- |
| $\tau_{00}$ item_no | 0.18 |
| $\tau_{11}$ subjnr.art_s_scale | 0.10 |
| $\tau_{11}$ item_no.art_s_scale | 0.02 |
| $\rho_{01}$ subjnr | -0.13 |
| $\rho_{01}$ item_no | 0.10 |
| ICC | 0.02 |
| $N_{\text{subjnr}}$ | 312 |
| $N_{\text{item\_no}}$ | 78 |
| Observations | 22108 |
| Marginal $R^2$ / Conditional $R^2$ | 0.120 / 0.141 |

**Appendix 2 – Table 8.** Model 7: N400_amp_mean ∼ noun_s_scale * iaf_scale + noun_s_scale * slope_scale + prestim_scale + (1+noun_s_scale|subjnr) + (1+noun_s_scale|item_no), data=combo_noun_sum

| *Predictors* | **N400_amp_mean** |  |  |  |
| --- | --- | --- | --- | --- |
|  | *Estimates* | *std. Error* | *CI (95%)* | *p* |
| (Intercept) | 1.13 | 0.12 | 0.91 – 1.36 | **<0.001** |
| noun s scale | -0.79 | 0.12 | -1.03 – -0.55 | **<0.001** |
| iaf scale | 0.13 | 0.07 | -0.01 – 0.27 | 0.070 |
| slope scale | -0.04 | 0.07 | -0.18 – 0.10 | 0.563 |
| prestim scale | -1.99 | 0.04 | -2.06 – -1.92 | **<0.001** |
| noun s scale × iaf scale | 0.03 | 0.04 | -0.05 – 0.10 | 0.473 |
| noun s scale × slope scale | -0.01 | 0.04 | -0.08 – 0.07 | 0.832 |
| **Random Effects** |  |  |  |  |
| $\sigma^2$ | 23.54 | | | |
| $\tau_{00}$ subjnr | 1.19 | | | |
| $\tau_{00}$ item_no | 0.49 | | | |
| $\tau_{11}$ subjnr.noun_s_scale | 0.06 | | | |
| $\tau_{11}$ item_no.noun_s_scale | 0.71 | | | |
| $\rho_{01}$ subjnr | 0.01 | | | |
| $\rho_{01}$ item_no | 0.33 | | | |
| ICC | 0.09 |  |  |  |
| N<sub>subjnr</sub> | 312 |  |  |  |
| N<sub>item_no</sub> | 78 |  |  |  |
| Observations | 19635 |  |  |  |
| Marginal R<sup>2</sup> / Conditional R<sup>2</sup> | 0.149 / 0.229 |  |  |  |

## Appendix 3: Outputs of SPP models with surprisal

**Appendix 3 – Table 1.** Model 1b: SPP_amp_mean ∼ cloze_scale * iaf_scale + cloze_scale * slope_scale + a1_s_scale + baseline_amp_scale + (1|subjnr) + (1|item_no), data= art_locs_sum

| *Predictors* | **SPP_amp_mean** |  |  |  |
| --- | --- | --- | --- | --- |
|  | *Estimates* | *std. Error* | *CI (95%)* | *p* |
| (Intercept) | -0.00 | 0.12 | -0.24 – 0.23 | 0.987 |
| cloze scale | 0.06 | 0.13 | -0.19 – 0.31 | 0.619 |
| iaf scale | 0.12 | 0.08 | -0.04 – 0.28 | 0.157 |
| slope scale | 0.03 | 0.08 | -0.13 – 0.19 | 0.696 |
| a1 s scale | 0.00 | 0.06 | -0.11 – 0.12 | 0.942 |
| baseline amp scale | -1.20 | 0.04 | -1.27 – -1.12 | **<0.001** |
| cloze scale × iaf scale | -0.03 | 0.08 | -0.19 – 0.13 | 0.748 |
| cloze scale × slope scale | -0.03 | 0.08 | -0.19 – 0.13 | 0.715 |
| **Random Effects** |  |  |  |  |
| $\sigma^2$ | 28.78 | | | |
| $\tau_{00}$ subjnr | 0.33 | | | |
| $\tau_{00}$ item_no | 0.16 | | | |
| ICC | 0.02 |  |  |  |
| $N_{\text{subjnr}}$ | 312 | | | |
| $N_{\text{item\_no}}$ | 80 | | | |
| Observations | 21922 |  |  |  |
| Marginal $R^2$ / Conditional $R^2$ | 0.046 / 0.062 | | | |

**Appendix 3 – Table 2.** Model 1a_surp: SPP_amp_mean ∼ art_s_scale * iaf_scale + art_s_scale * slope_scale + baseline_amp_scale + (1|subjnr) + (1|item_no), data= art_locs_sum

| *Predictors* | **SPP_amp_mean** |  |  |  |
| --- | --- | --- | --- | --- |
|  | *Estimates* | *std. Error* | *CI (95%)* | *p* |
| (Intercept) | 0.05 | 0.07 | -0.08 – 0.18 | 0.494 |
| art s scale | -0.03 | 0.06 | -0.16 – 0.09 | 0.590 |
| iaf scale | 0.10 | 0.05 | -0.00 – 0.20 | 0.054 |
| slope scale | 0.01 | 0.05 | -0.09 – 0.11 | 0.875 |
| baseline amp scale | -1.20 | 0.04 | -1.27 – -1.12 | **<0.001** |
| art s scale × iaf scale | 0.03 | 0.04 | -0.05 – 0.12 | 0.404 |
| art s scale × slope scale | -0.01 | 0.04 | -0.09 – 0.07 | 0.843 |
| **Random Effects** |  |  |  |  |
| $\sigma^2$ | 28.78 | | | |
| $\tau_{00}$ subjnr | 0.33 | | | |
| $\tau_{00}$ item_no | 0.16 | | | |
| ICC | 0.02 |  |  |  |
| N<sub>subjnr</sub> | 312 |  |  |  |
| N<sub>item_no</sub> | 80 |  |  |  |
| Observations | 21922 |  |  |  |
| Marginal R<sup>2</sup> / Conditional R<sup>2</sup> | 0.046 / 0.062 |  |  |  |

**Appendix 3 – Table 3.**
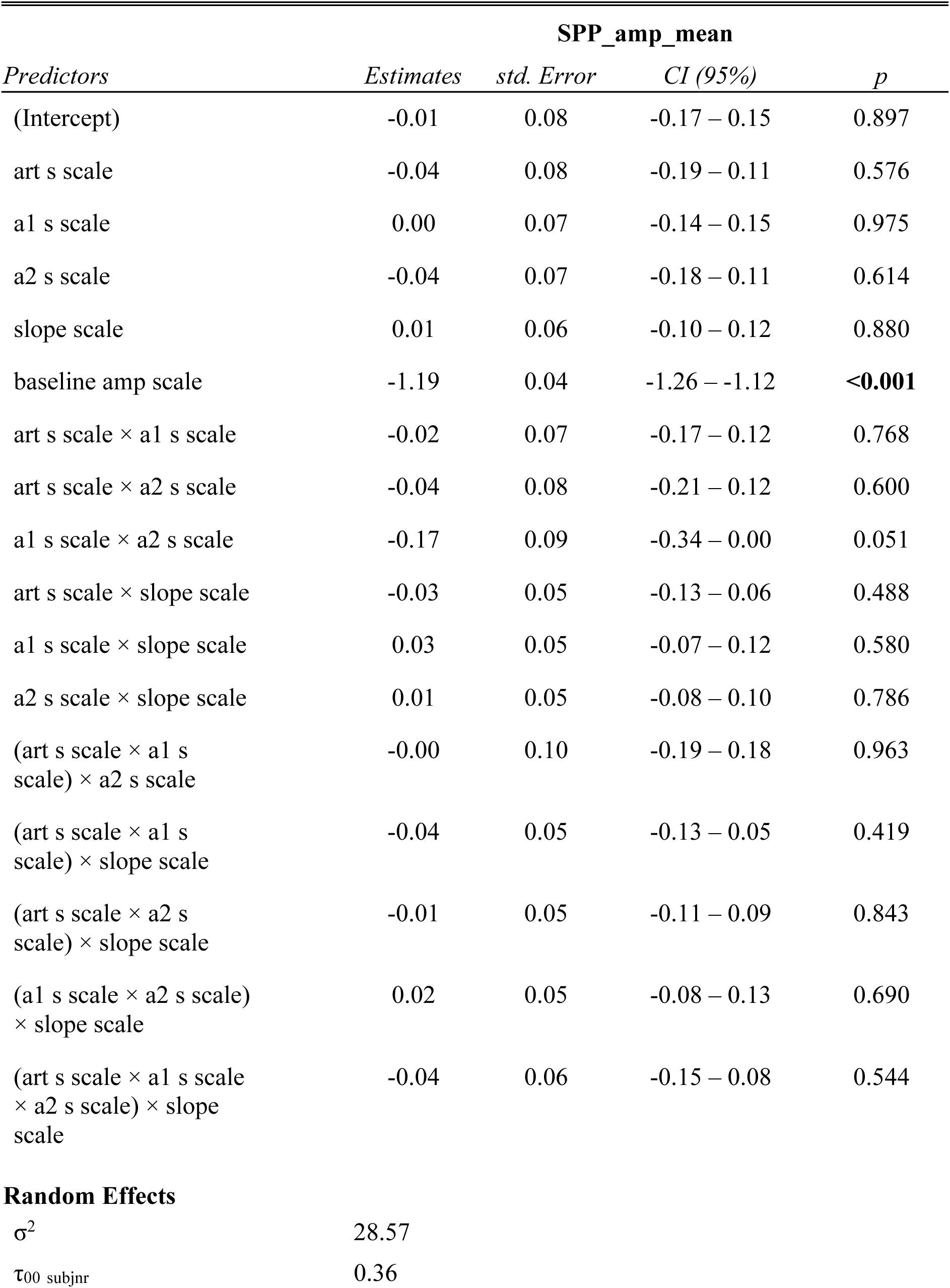
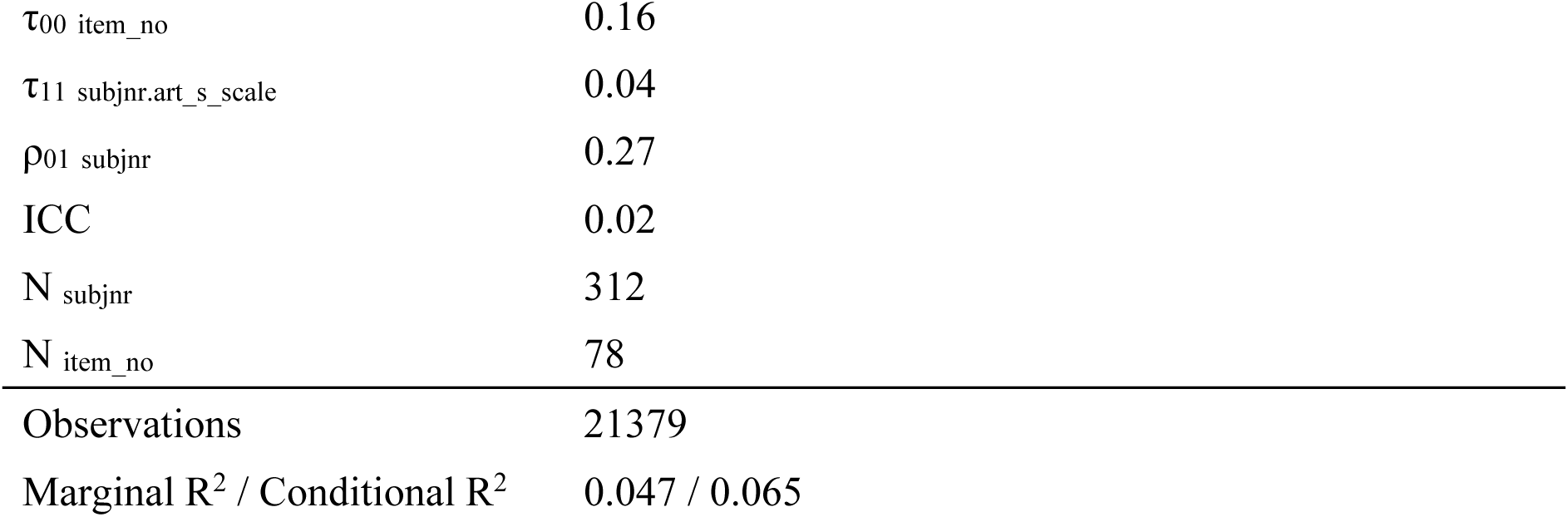
Model 1c_slope: SPP_amp_mean ∼ art_s_scale * a1_s_scale * a2_s_scale * slope_scale + baseline_amp_scale + (1+art_s_scale|subjnr) + (1|item_no), data= art_locs_sum

| *Predictors* | **SPP_amp_mean** |  |  |  |
| --- | --- | --- | --- | --- |
|  | *Estimates* | *std. Error* | *CI (95%)* | *p* |
| (Intercept) | -0.01 | 0.08 | -0.17 – 0.15 | 0.897 |
| art s scale | -0.04 | 0.08 | -0.19 – 0.11 | 0.576 |
| a1 s scale | 0.00 | 0.07 | -0.14 – 0.15 | 0.975 |
| a2 s scale | -0.04 | 0.07 | -0.18 – 0.11 | 0.614 |
| slope scale | 0.01 | 0.06 | -0.10 – 0.12 | 0.880 |
| baseline amp scale | -1.19 | 0.04 | -1.26 – -1.12 | **<0.001** |
| art s scale × a1 s scale | -0.02 | 0.07 | -0.17 – 0.12 | 0.768 |
| art s scale × a2 s scale | -0.04 | 0.08 | -0.21 – 0.12 | 0.600 |
| a1 s scale × a2 s scale | -0.17 | 0.09 | -0.34 – 0.00 | 0.051 |
| art s scale × slope scale | -0.03 | 0.05 | -0.13 – 0.06 | 0.488 |
| a1 s scale × slope scale | 0.03 | 0.05 | -0.07 – 0.12 | 0.580 |
| a2 s scale × slope scale | 0.01 | 0.05 | -0.08 – 0.10 | 0.786 |
| (art s scale × a1 s scale) × a2 s scale | -0.00 | 0.10 | -0.19 – 0.18 | 0.963 |
| (art s scale × a1 s scale) × slope scale | -0.04 | 0.05 | -0.13 – 0.05 | 0.419 |
| (art s scale × a2 s scale) × slope scale | -0.01 | 0.05 | -0.11 – 0.09 | 0.843 |
| (a1 s scale × a2 s scale) × slope scale | 0.02 | 0.05 | -0.08 – 0.13 | 0.690 |
| (art s scale × a1 s scale × a2 s scale) × slope scale | -0.04 | 0.06 | -0.15 – 0.08 | 0.544 |
| **Random Effects** |  |  |  |  |
| $\sigma^2$ | 28.57 | | | |
| $\tau_{00}$ subjnr | 0.36 | | | |

|  |  |
| --- | --- |
| $\tau_{00}$ item_no | 0.16 |
| $\tau_{11}$ subjnr.art_s_scale | 0.04 |
| $\rho_{01}$ subjnr | 0.27 |
| ICC | 0.02 |
| $N_{\text{subjnr}}$ | 312 |
| $N_{\text{item\_no}}$ | 78 |
| Observations | 21379 |
| Marginal $R^2$ / Conditional $R^2$ | 0.047 / 0.065 |

**Appendix 3 – Table 4.**
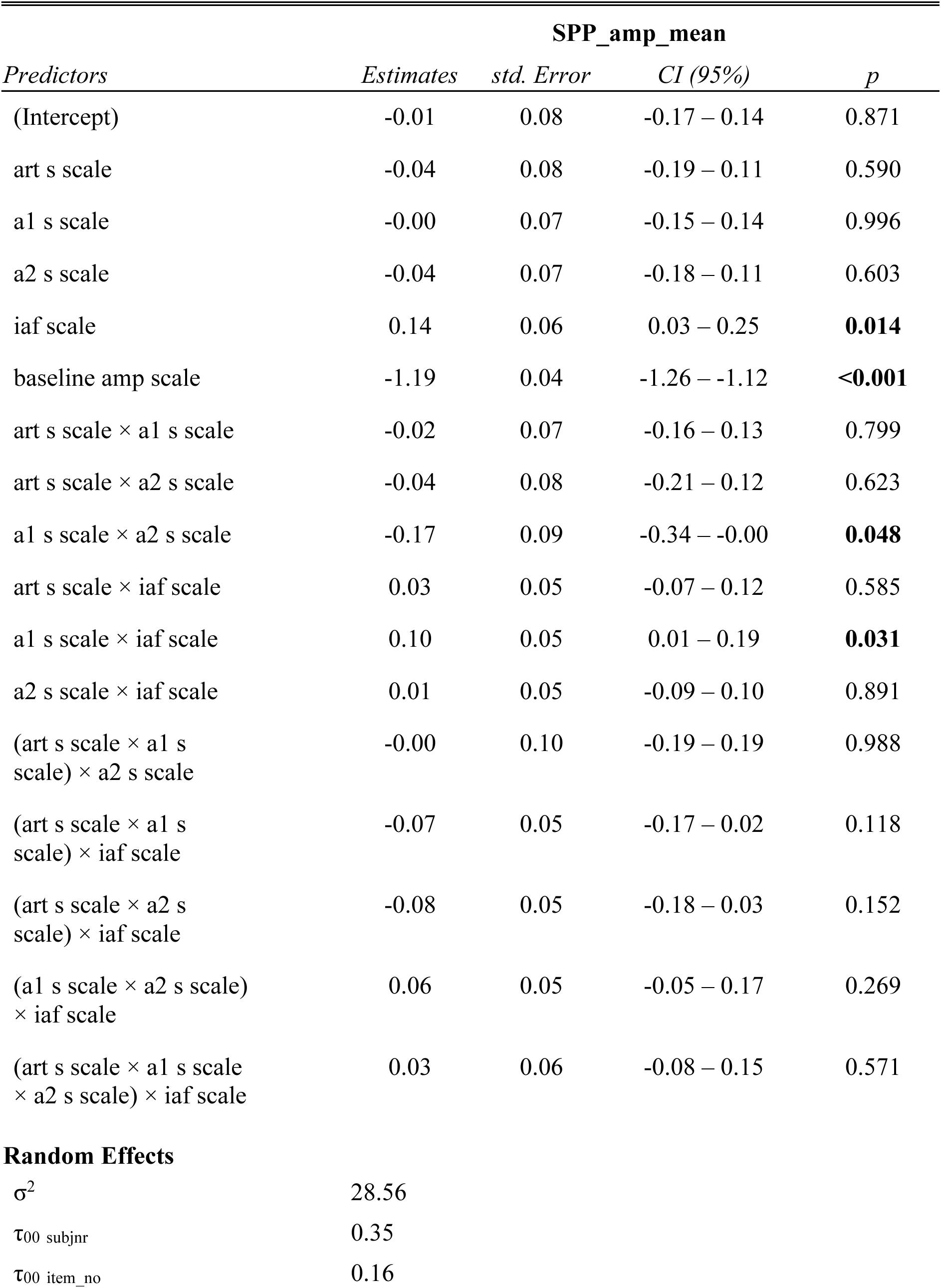
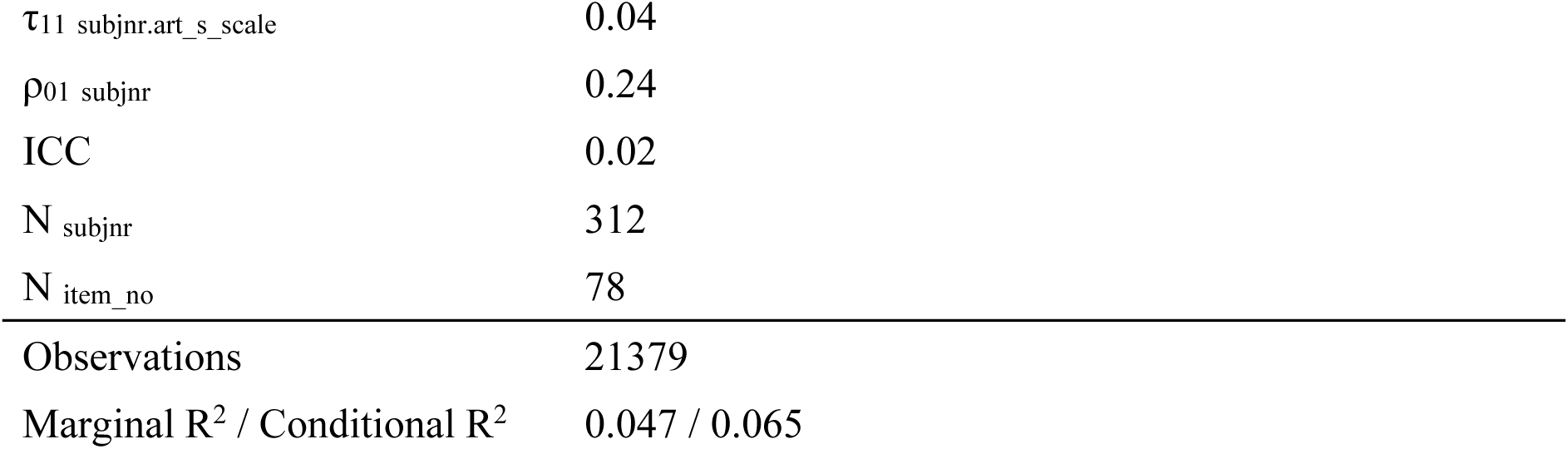
Model 1c_iaf: SPP_amp_mean ∼ art_s_scale * a1_s_scale * a2_s_scale * slope_scale + baseline_amp_scale + (1+art_s_scale|subjnr) + (1|item_no), data= art_locs_sum

| *Predictors* | **SPP_amp_mean** |  |  |  |
| --- | --- | --- | --- | --- |
|  | *Estimates* | *std. Error* | *CI (95%)* | *p* |
| (Intercept) | -0.01 | 0.08 | -0.17 – 0.14 | 0.871 |
| art s scale | -0.04 | 0.08 | -0.19 – 0.11 | 0.590 |
| a1 s scale | -0.00 | 0.07 | -0.15 – 0.14 | 0.996 |
| a2 s scale | -0.04 | 0.07 | -0.18 – 0.11 | 0.603 |
| iaf scale | 0.14 | 0.06 | 0.03 – 0.25 | **0.014** |
| baseline amp scale | -1.19 | 0.04 | -1.26 – -1.12 | **<0.001** |
| art s scale × a1 s scale | -0.02 | 0.07 | -0.16 – 0.13 | 0.799 |
| art s scale × a2 s scale | -0.04 | 0.08 | -0.21 – 0.12 | 0.623 |
| a1 s scale × a2 s scale | -0.17 | 0.09 | -0.34 – -0.00 | **0.048** |
| art s scale × iaf scale | 0.03 | 0.05 | -0.07 – 0.12 | 0.585 |
| a1 s scale × iaf scale | 0.10 | 0.05 | 0.01 – 0.19 | **0.031** |
| a2 s scale × iaf scale | 0.01 | 0.05 | -0.09 – 0.10 | 0.891 |
| (art s scale × a1 s scale) × a2 s scale | -0.00 | 0.10 | -0.19 – 0.19 | 0.988 |
| (art s scale × a1 s scale) × iaf scale | -0.07 | 0.05 | -0.17 – 0.02 | 0.118 |
| (art s scale × a2 s scale) × iaf scale | -0.08 | 0.05 | -0.18 – 0.03 | 0.152 |
| (a1 s scale × a2 s scale) × iaf scale | 0.06 | 0.05 | -0.05 – 0.17 | 0.269 |
| (art s scale × a1 s scale × a2 s scale) × iaf scale | 0.03 | 0.06 | -0.08 – 0.15 | 0.571 |
| **Random Effects** |  |  |  |  |
| $\sigma^2$ | 28.56 | | | |
| $\tau_{00}$ subjnr | 0.35 | | | |
| $\tau_{00}$ item_no | 0.16 | | | |

|  |  |
| --- | --- |
| $\tau_{11}$ subjnr.art_s_scale | 0.04 |
| $\rho_{01}$ subjnr | 0.24 |
| ICC | 0.02 |
| $N_{\text{subjnr}}$ | 312 |
| $N_{\text{item\_no}}$ | 78 |
| Observations | 21379 |
| Marginal $R^2$ / Conditional $R^2$ | 0.047 / 0.065 |

**Appendix 3 – Table 5.** Model 2b: SPP_amp_mean ∼ cloze_scale * iaf_scale + cloze_scale * slope_scale + baseline_amp_scale + art_s_scale + (1|subjnr) + (1+cloze_scale|item_no), data= noun_locs_sum

| *Predictors* | **SPP_amp_mean** |  |  |  |
| --- | --- | --- | --- | --- |
|  | *Estimates* | *std. Error* | *CI (95%)* | *p* |
| (Intercept) | -1.04 | 0.07 | -1.18 – -0.90 | **<0.001** |
| cloze scale | 0.11 | 0.05 | 0.01 – 0.20 | **0.030** |
| iaf scale | -0.01 | 0.05 | -0.11 – 0.10 | 0.872 |
| slope scale | 0.04 | 0.05 | -0.06 – 0.14 | 0.450 |
| baseline amp scale | -0.54 | 0.04 | -0.61 – -0.46 | **<0.001** |
| art s scale | 0.03 | 0.05 | -0.07 – 0.14 | 0.558 |
| cloze scale × iaf scale | -0.05 | 0.04 | -0.12 – 0.02 | 0.199 |
| cloze scale × slope scale | -0.00 | 0.04 | -0.07 – 0.07 | 0.974 |
| **Random Effects** |  |  |  |  |
| $\sigma^2$ | 27.66 | | | |
| $\tau_{00}$ subjnr | 0.43 | | | |
| $\tau_{00}$ item_no | 0.17 | | | |
| $\tau_{11}$ item_no.cloze_scale | 0.08 | | | |
| $\rho_{01}$ item_no | -0.18 | | | |
| ICC | 0.02 |  |  |  |
| $N_{\text{subjnr}}$ | 312 | | | |
| $N_{\text{item\_no}}$ | 80 | | | |
| Observations | 21282 |  |  |  |
| Marginal R<sup>2</sup> / Conditional R<sup>2</sup> | 0.011 / 0.035 |  |  |  |

**Appendix 3 – Table 6.**
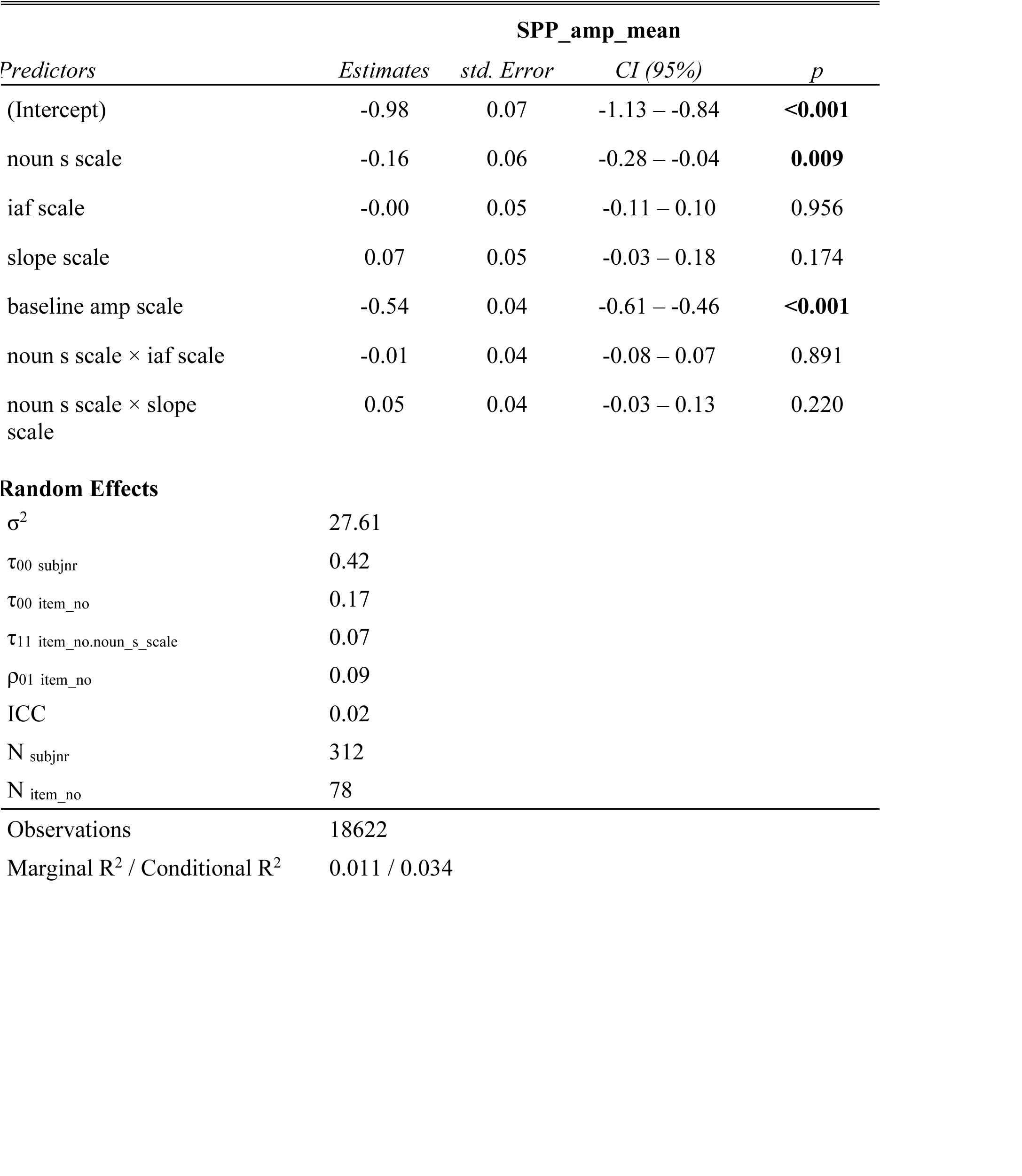
Model 2a_surp: SPP_amp_mean ∼ noun_s_scale * iaf_scale + noun_s_scale * slope_scale + baseline_amp_scale + (1|subjnr) + (1+noun_s_scale|item_no), data= noun_locs_sum

| *Predictors* | **SPP_amp_mean** |  |  |  |
| --- | --- | --- | --- | --- |
|  | *Estimates* | *std. Error* | *CI (95%)* | *p* |
| (Intercept) | -0.98 | 0.07 | -1.13 – -0.84 | **<0.001** |
| noun s scale | -0.16 | 0.06 | -0.28 – -0.04 | **0.009** |
| iaf scale | -0.00 | 0.05 | -0.11 – 0.10 | 0.956 |
| slope scale | 0.07 | 0.05 | -0.03 – 0.18 | 0.174 |
| baseline amp scale | -0.54 | 0.04 | -0.61 – -0.46 | **<0.001** |
| noun s scale × iaf scale | -0.01 | 0.04 | -0.08 – 0.07 | 0.891 |
| noun s scale × slope scale | 0.05 | 0.04 | -0.03 – 0.13 | 0.220 |
| **Random Effects** |  |  |  |  |
| $\sigma^2$ | 27.61 | | | |
| $\tau_{00}$ subjnr | 0.42 | | | |
| $\tau_{00}$ item_no | 0.17 | | | |
| $\tau_{11}$ item_no.noun_s_scale | 0.07 | | | |
| $\rho_{01}$ item_no | 0.09 | | | |
| ICC | 0.02 |  |  |  |
| N<sub>subjnr</sub> | 312 |  |  |  |
| N<sub>item_no</sub> | 78 |  |  |  |
| Observations | 18622 |  |  |  |
| Marginal R<sup>2</sup> / Conditional R<sup>2</sup> | 0.011 / 0.034 |  |  |  |

**Appendix 3 – Table 7.**
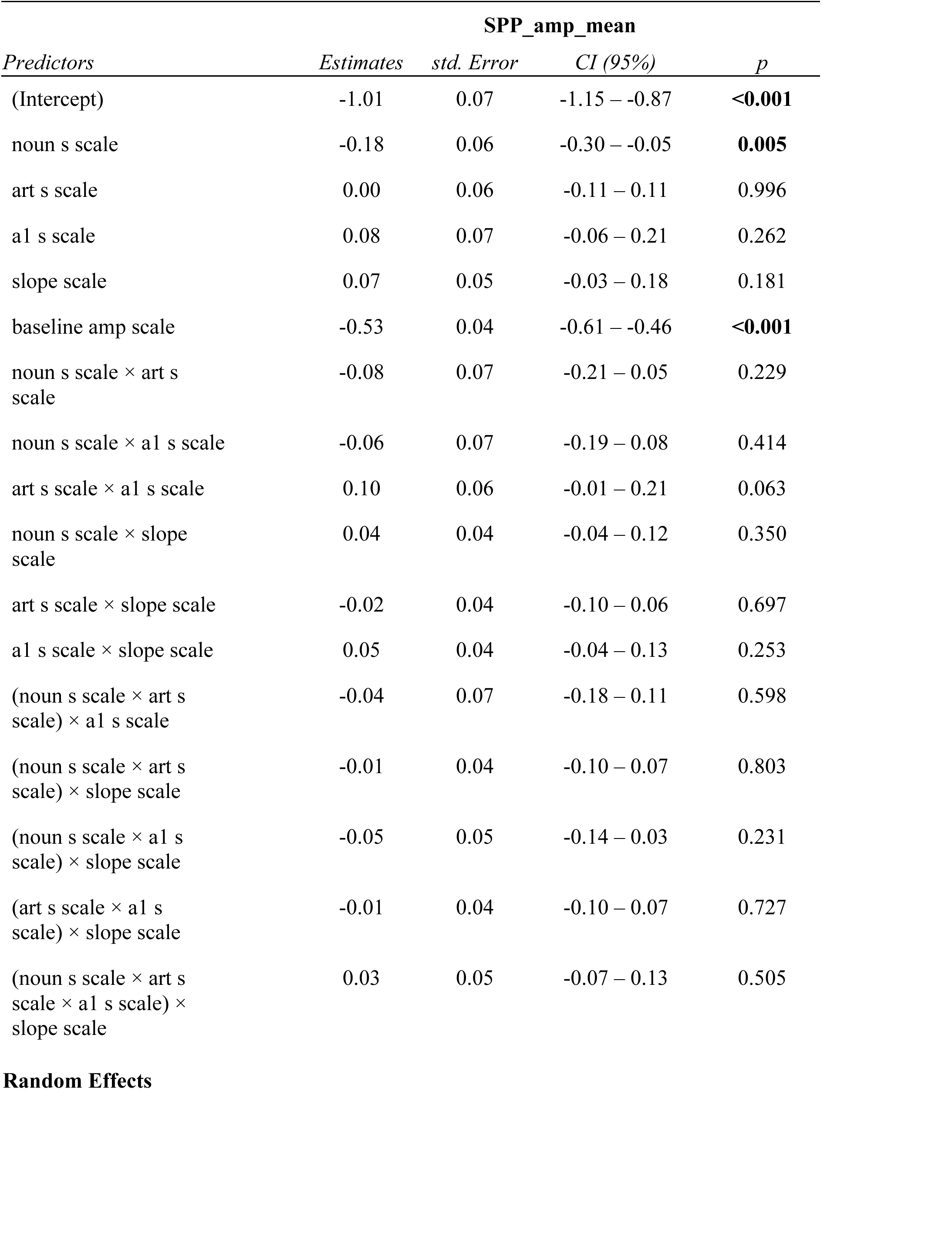
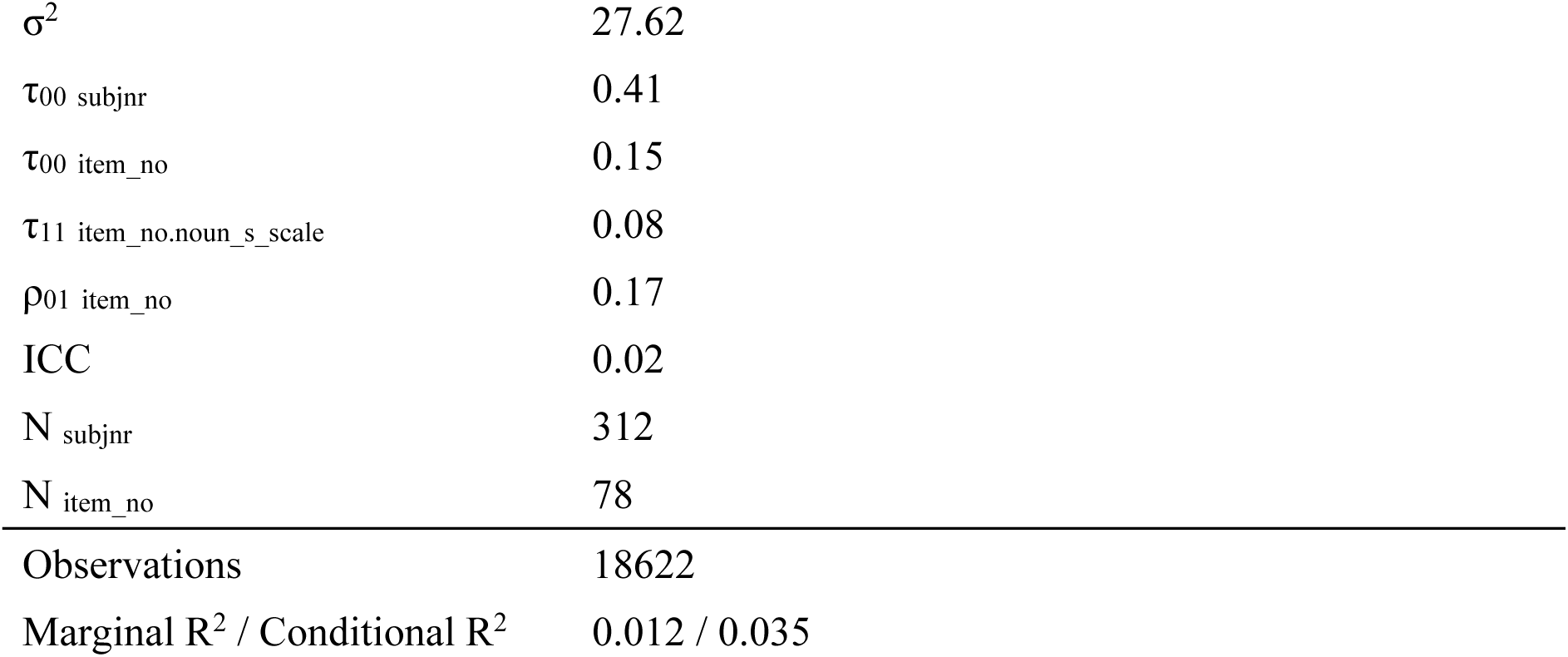
Model 2c_slope: SPP_amp_mean ∼ noun_s_scale * art_s_scale * a1_s_scale * slope_scale + baseline_amp_scale + (1|subjnr) + (1+noun_s_scale|item_no), data= noun_locs_sum

| *Predictors* | **SPP_amp_mean** |  |  |  |
| --- | --- | --- | --- | --- |
|  | *Estimates* | *std. Error* | *CI (95%)* | *p* |
| (Intercept) | -1.01 | 0.07 | -1.15 – -0.87 | **<0.001** |
| noun s scale | -0.18 | 0.06 | -0.30 – -0.05 | **0.005** |
| art s scale | 0.00 | 0.06 | -0.11 – 0.11 | 0.996 |
| a1 s scale | 0.08 | 0.07 | -0.06 – 0.21 | 0.262 |
| slope scale | 0.07 | 0.05 | -0.03 – 0.18 | 0.181 |
| baseline amp scale | -0.53 | 0.04 | -0.61 – -0.46 | **<0.001** |
| noun s scale × art s scale | -0.08 | 0.07 | -0.21 – 0.05 | 0.229 |
| noun s scale × a1 s scale | -0.06 | 0.07 | -0.19 – 0.08 | 0.414 |
| art s scale × a1 s scale | 0.10 | 0.06 | -0.01 – 0.21 | 0.063 |
| noun s scale × slope scale | 0.04 | 0.04 | -0.04 – 0.12 | 0.350 |
| art s scale × slope scale | -0.02 | 0.04 | -0.10 – 0.06 | 0.697 |
| a1 s scale × slope scale | 0.05 | 0.04 | -0.04 – 0.13 | 0.253 |
| (noun s scale × art s scale) × a1 s scale | -0.04 | 0.07 | -0.18 – 0.11 | 0.598 |
| (noun s scale × art s scale) × slope scale | -0.01 | 0.04 | -0.10 – 0.07 | 0.803 |
| (noun s scale × a1 s scale) × slope scale | -0.05 | 0.05 | -0.14 – 0.03 | 0.231 |
| (art s scale × a1 s scale) × slope scale | -0.01 | 0.04 | -0.10 – 0.07 | 0.727 |
| (noun s scale × art s scale × a1 s scale) × slope scale | 0.03 | 0.05 | -0.07 – 0.13 | 0.505 |

Random Effects
|  |  |
| --- | --- |
| $\sigma^2$ | 27.62 |
| $\tau_{00}$ subjnr | 0.41 |
| $\tau_{00}$ item_no | 0.15 |
| $\tau_{11}$ item_no.noun_s_scale | 0.08 |
| $\rho_{01}$ item_no | 0.17 |
| ICC | 0.02 |
| N subjnr | 312 |
| N item_no | 78 |
| Observations | 18622 |
| Marginal R<sup>2</sup> / Conditional R<sup>2</sup> | 0.012 / 0.035 |

**Appendix 3 – Table 8.**
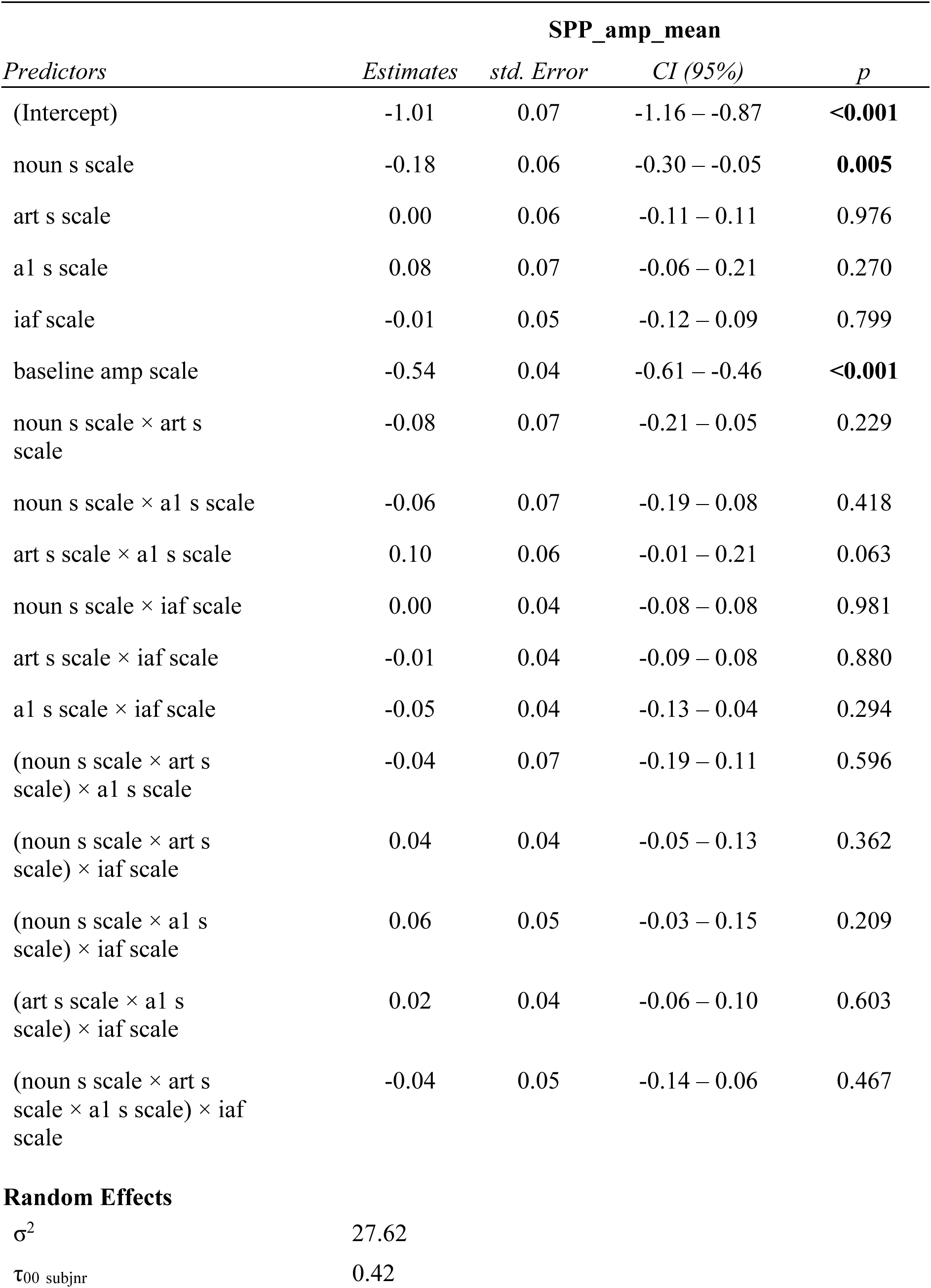
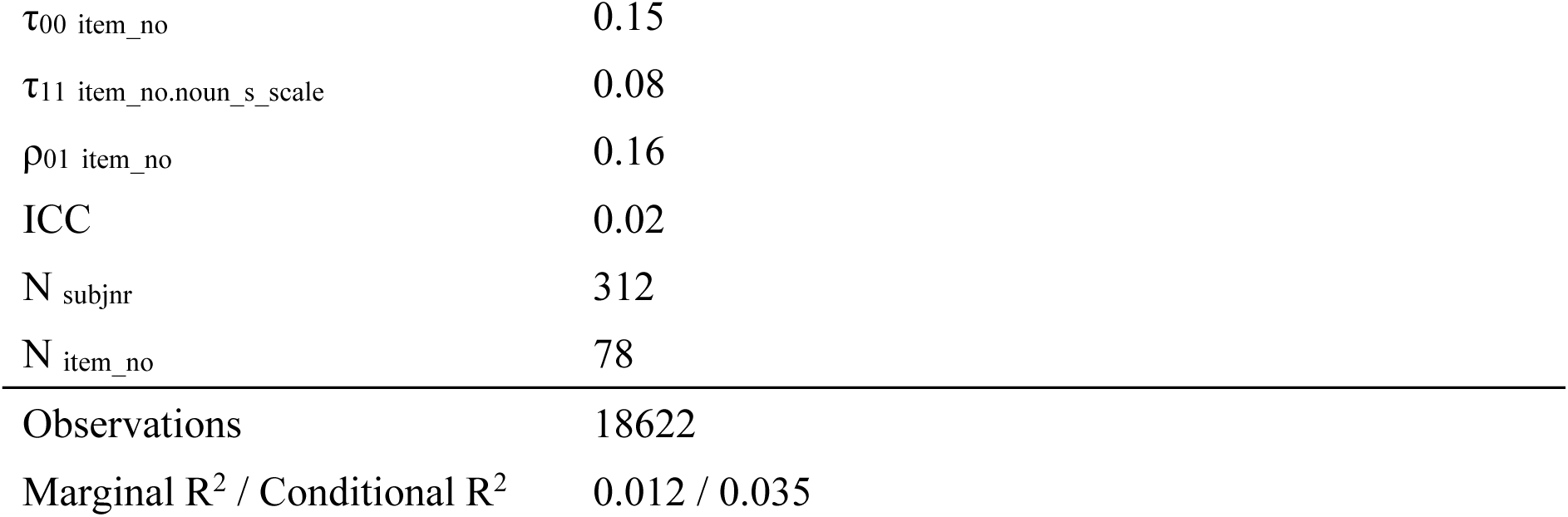
Model 2c_iaf: SPP_amp_mean ∼ noun_s_scale * art_s_scale * a1_s_scale * iaf_scale + baseline_amp_scale + (1|subjnr) + (1+noun_s_scale|item_no), data= noun_locs_sum

| *Predictors* | **SPP_amp_mean** |  |  |  |
| --- | --- | --- | --- | --- |
|  | *Estimates* | *std. Error* | *CI (95%)* | *p* |
| (Intercept) | -1.01 | 0.07 | -1.16 – -0.87 | **<0.001** |
| noun s scale | -0.18 | 0.06 | -0.30 – -0.05 | **0.005** |
| art s scale | 0.00 | 0.06 | -0.11 – 0.11 | 0.976 |
| a1 s scale | 0.08 | 0.07 | -0.06 – 0.21 | 0.270 |
| iaf scale | -0.01 | 0.05 | -0.12 – 0.09 | 0.799 |
| baseline amp scale | -0.54 | 0.04 | -0.61 – -0.46 | **<0.001** |
| noun s scale × art s scale | -0.08 | 0.07 | -0.21 – 0.05 | 0.229 |
| noun s scale × a1 s scale | -0.06 | 0.07 | -0.19 – 0.08 | 0.418 |
| art s scale × a1 s scale | 0.10 | 0.06 | -0.01 – 0.21 | 0.063 |
| noun s scale × iaf scale | 0.00 | 0.04 | -0.08 – 0.08 | 0.981 |
| art s scale × iaf scale | -0.01 | 0.04 | -0.09 – 0.08 | 0.880 |
| a1 s scale × iaf scale | -0.05 | 0.04 | -0.13 – 0.04 | 0.294 |
| (noun s scale × art s scale) × a1 s scale | -0.04 | 0.07 | -0.19 – 0.11 | 0.596 |
| (noun s scale × art s scale) × iaf scale | 0.04 | 0.04 | -0.05 – 0.13 | 0.362 |
| (noun s scale × a1 s scale) × iaf scale | 0.06 | 0.05 | -0.03 – 0.15 | 0.209 |
| (art s scale × a1 s scale) × iaf scale | 0.02 | 0.04 | -0.06 – 0.10 | 0.603 |
| (noun s scale × art s scale × a1 s scale) × iaf scale | -0.04 | 0.05 | -0.14 – 0.06 | 0.467 |
| **Random Effects** |  |  |  |  |
| $\sigma^2$ | 27.62 | | | |
| $\tau_{00}$ subjnr | 0.42 | | | |

|  |  |
| --- | --- |
| $\tau_{00}$ item_no | 0.15 |
| $\tau_{11}$ item_no.noun_s_scale | 0.08 |
| $\rho_{01}$ item_no | 0.16 |
| ICC | 0.02 |
| $N_{\text{subjnr}}$ | 312 |
| $N_{\text{item\_no}}$ | 78 |
| Observations | 18622 |
| Marginal $R^2$ / Conditional $R^2$ | 0.012 / 0.035 |

## Appendix 4 Exploratory Bayesian sensitivity analyses

To further quantify the magnitude of the effect of cloze probability on SPP amplitude, two Bayesian sensitivity analyses were conducted using the *brms* package in *R*, which is based on the Stan programming language (Bürkner, 2017). The first model tested the effect of article cloze on pre-article SPP amplitude, while the second model tested the effect of noun cloze on pre-noun SPP amplitude. The basic model structure of the models of interest (M_1_) is presented below:

`SPP_amp_mean ∼ c_cloze + baseline_amp_scale + (c_cloze|subjnr) + (c_cloze|item_no)`

These models were compared against corresponding null models (M_0_), which took the following form:

`SPP_amp_mean ∼ 1 + baseline_amp_scale + (c_cloze|subjnr) + (c_cloze|item_no)`

We adopted a similar strategy as detailed in Nicenboim et al. (2024; Chapter 15). Cloze probability was mean-centered, and priors for predictors (*β*) were assigned to a normal distribution with a mean of zero and with a range of standard deviations (from 0.5 to 10). This allowed for the manipulation of the amount of expected variance around an effect, facilitating the computation of multiple Bayes factors. The settings for the remaining priors were based on Nicenboim et al.’s (2020) uninformative priors. This decision was based on the limited availability of information from existing SPP studies, which predominantly employ t-tests or ANOVAs. The specifications for priors not relating to predictors (*β*) in *brms* for all models were as follows:

`other_priors <-c ( prior(normal(0, 10), class = Intercept), prior(normal(0, 10), class = sigma), prior(normal(0, 10), class = sd), prior(lkj(2), class = cor))`

Sensitivity analyses yield Bayes factors indicating the amount of evidence for or against the effects, with a Bayes factor below one (BF_10_ < 1) representing evidence in favour of M_0_, and a Bayes factor above one (BF_10_ > 1) reflecting evidence for M_1_ (Nicenboim et al., 2024; Chapter 15).

### Results

#### Article results

The analysis at the article yielded divergent transitions, which may be reflective of poor model fit. To combat this, the priors were adjusted based on Nicenboim et al. (2024; see below). However, these priors also led to divergent transitions. Therefore, whilst we present the results of the sensitivity analysis with the uninformative priors here, it is recommended that such results be interpreted with caution (See *Figure 1*).

`priors1 <-c(`

`prior(normal(2, 5), class = Intercept), prior(normal(10, 5), class = sigma), prior(normal(0, 2), class = sd), prior(lkj(4), class = cor))`

#### Noun results

In contrast to the article results, the sensitivity analysis at the noun with the uninformative priors ran successfully, revealing Bayes factors below 1 for all prior standard deviations (See *Figure 1* below). This suggests stronger evidence for the null model (M_0_) over M_1_, providing evidence against an effect of noun cloze on pre-noun SPP amplitude.

**Figure 1.**
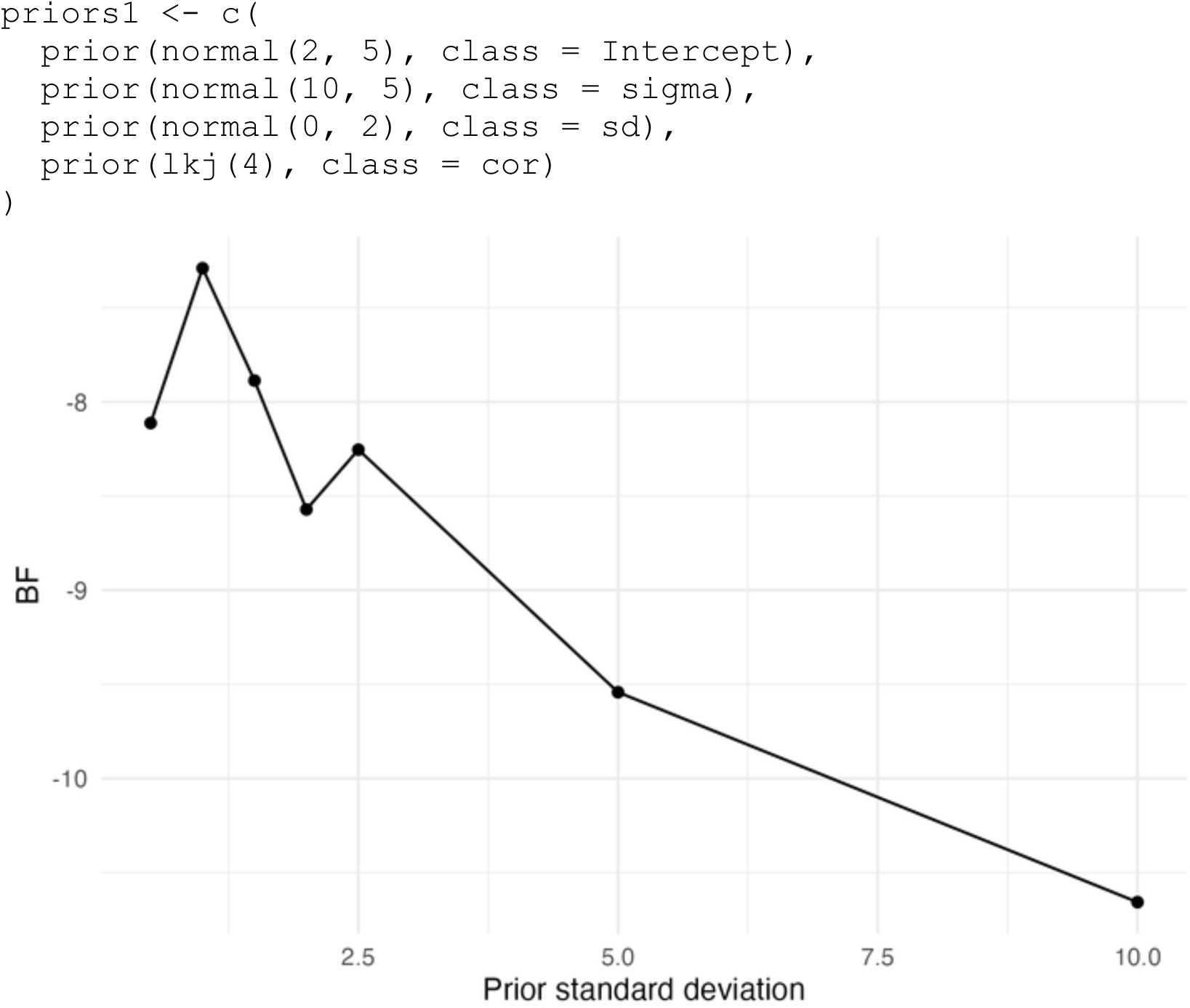
The number of participants at each stage of the data analysis, including the number removed and reasons for removal.

**Figure 2.**
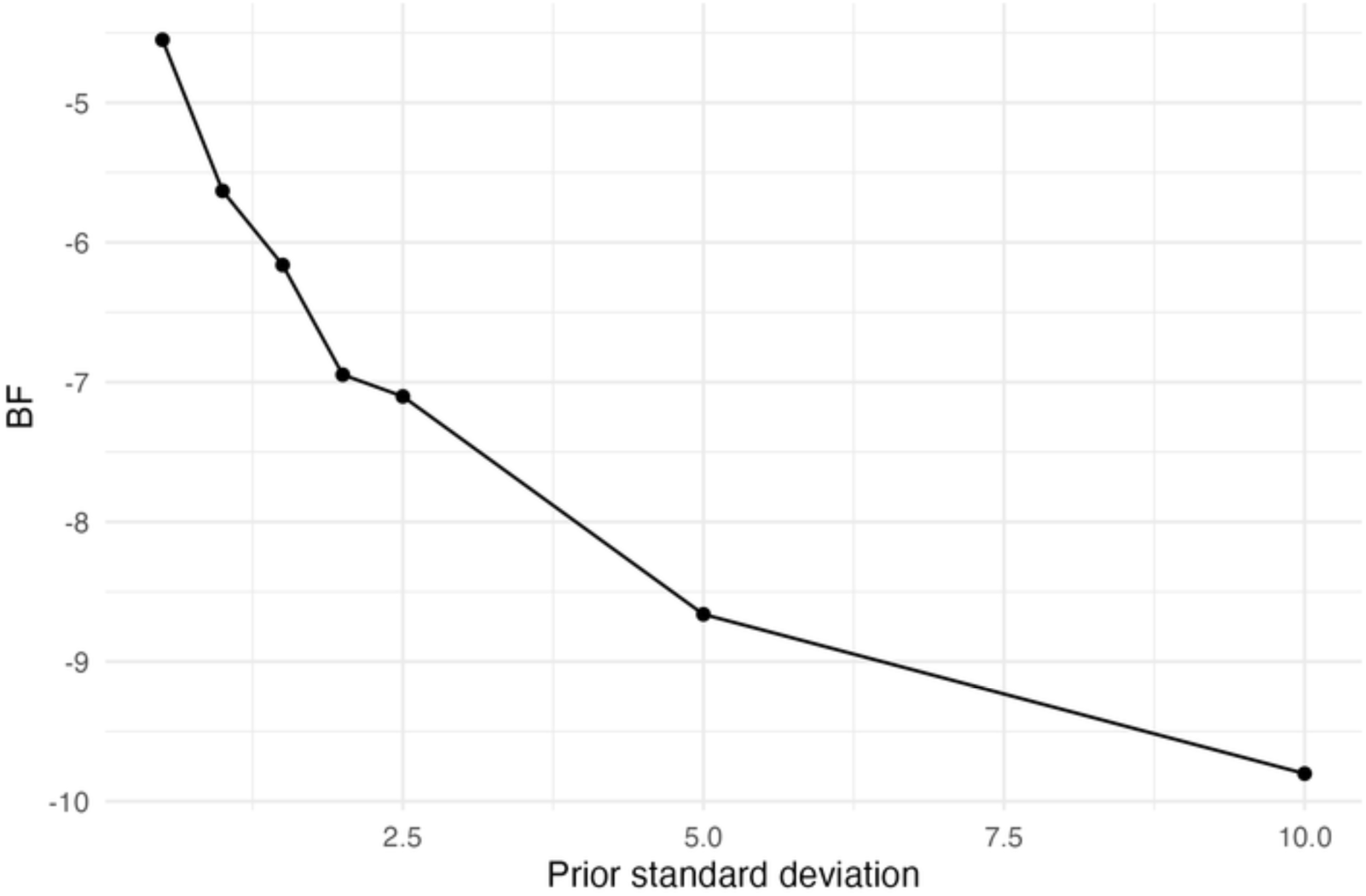
SPP ERP grand average plots for articles over channels of interest according to between-sentence article cloze probability (*n* = 312) to gauge SPP effects. Blue lines represent high article cloze probability (>50%) whilst orange dashed lines reflect low article cloze probability (<50%). Blue and orange shading reflects the standard errors for each condition. The SPP window is shaded grey, and the black dashed line depicts the onset of the article word. Note that the plot depicts non-baseline corrected ERP activity.

